# Interpretations of microbial community studies are biased by the selected 16S rRNA gene amplicon sequencing pipeline

**DOI:** 10.1101/2019.12.17.880468

**Authors:** Daniel Straub, Nia Blackwell, Adrian Langarica Fuentes, Alexander Peltzer, Sven Nahnsen, Sara Kleindienst

## Abstract

One of the major methods to identify microbial community composition, to unravel microbial population dynamics, and to explore microbial diversity in environmental samples is DNA- or RNA-based 16S rRNA (gene) amplicon sequencing. Subsequent bioinformatics analyses are required to extract valuable information from the high-throughput sequencing approach. However, manifold bioinformatics tools complicate their choice and might cause differences in data interpretation, making the selection of the pipeline a crucial step.

Here, we compared the performance of most widely used 16S rRNA gene amplicon sequencing analysis tools (i.e. Mothur, QIIME1, QIIME2, and MEGAN) using mock datasets and environmental samples from contrasting terrestrial and freshwater sites. Our results showed that QIIME2 outcompeted all other investigated tools in sequence recovery (>10 times less false positives), taxonomic assignments (>22% better F-score) and diversity estimates (>5% better assessment), while there was still room for improvement e.g. imperfect sequence recovery (recall up to 87%) or detection of additional false sequences (precision up to 72%). Furthermore, we found that microbial diversity estimates and highest abundant taxa varied among analysis pipelines (i.e. only one in five genera was shared among all analysis tools) when analyzing environmental samples, which might skew biological conclusions.

Our findings were subsequently implemented in a high-performance computing conformant workflow following the FAIR (Findable, Accessible, Interoperable, and Re-usable) principle, allowing reproducible 16S rRNA gene amplicon sequence analysis starting from raw sequence files. Our presented workflow can be utilized for future studies, thereby facilitating the analysis of high-throughput DNA- or RNA-based 16S rRNA (gene) sequencing data substantially.

**Importance:** Microorganisms play an essential role in biogeochemical cycling events across the globe. Phylogenetic marker gene analysis is a widely used method to explore microbial community dynamics in space and time, to predict the ecological relevance of microbial populations, or to identify microbial key players in biogeochemical cycles. Several computational analysis methods were developed to aid 16S rRNA gene analysis but choosing the best method is not trivial. In this study, we compared popular analysis methods (i.e. Mothur, QIIME1 and 2, and MEGAN) using samples with known microbial composition (i.e. mock community samples) and environmental samples from contrasting habitats (i.e. groundwater, soil, sediment, and river water). Our findings provide guidance for choosing the currently optimal 16S rRNA gene sequencing analysis method and we implemented our recommended pipeline into a reproducible workflow, which follows highest bioinformatics standards and is open source and free to use.

## 1 Introduction

The ribosomal 16S rRNA gene is a phylogenetic marker that has been analyzed extensively within the last decade due to its presence in all microorganisms (Hugenholtz, Goebel et al. 1998), and due to a combination of variable regions, influenced by the evolutionary clock that allow differentiation of taxa, with conserved regions, for universal priming (Head, Saunders et al. 1998). Since the dawn of second-generation sequencing methods, the cost of nucleotide sequencing has decreased dramatically (Wetterstrand 2018) and DNA- or RNA-based 16S rRNA (gene) amplicon sequencing is becoming more and more affordable. Initially, 454 pyrosequencing was employed but, after resolving early limitations, relatively short Illumina sequencing is currently dominating (Claesson, Wang et al. 2010, D’Amore, Ijaz et al. 2016) because of higher sequence quality and cost advantages. Next-generation sequencing (NGS) is able to produce a vast number of sequences at unprecedented speed. Data processing and analysis require significant computational power (Muir, Li et al. 2016) and are rapidly becoming the bottleneck in NGS experiments. Compared to other sequencing strategies such as whole-genome sequencing of large genomes, 16S rRNA gene amplicon sequencing is relatively easy on computational requirements, however, these are increasing with the number of samples, sequencing depth, community complexity and size of the taxonomic reference database (Almeida, Mitchell et al. 2018). Additionally, analysis methods and employed software influence the required computational resources (Nearing, Douglas et al. 2018).

16S rRNA gene amplicon sequencing analysis pipelines are required to be user-friendly and to provide the best output possible. Criteria for optimal results include the recovery of all 16S rRNA gene amplicon sequences and taxa (full sensitivity) with no false positive detection (full specificity). Also, *in situ* relative abundances are ideally perfectly represented. However, all current analysis methods suffer from imperfect recall (not all sequences or taxa are detected) or imperfect precision (additional false sequences or taxa are detected) (Callahan, McMurdie et al. 2016) that originate from a diverse set of frequent shortcomings of the entire workflow. These include biases in sample preparation (e.g. DNA extraction, PCR, sequencing library preparation), suboptimal experimental design (e.g. amplicon selection), erroneous sequences produced by the sequencing method and the bioinformatics analysis strategy (Kozich, Westcott et al. 2013, Wesolowska-Andersen, Bahl et al. 2014, de Muinck, Trosvik et al. 2017, Laursen, Dalgaard et al. 2017, Almeida, Mitchell et al. 2018, Nearing, Douglas et al. 2018).

The scientific literature suggesting software applications for the analysis of 16S rRNA gene sequencing data is continuously growing and many pipelines have been proposed. Among the most prominent tools to date are QIIME (Caporaso, Kuczynski et al. 2010), Mothur (Schloss, Westcott et al. 2009) and MEGAN (Mitra, Stärk et al. 2011). With each amassing around ten thousand citations, Mothur (Schloss, Westcott et al. 2009) and QIIME1 (Caporaso, Kuczynski et al. 2010) were the most popular tools for amplicon sequencing analysis in recent years. Both tools cluster sequences with a given similarity (e.g., >=97%) into operational taxonomic units (OTUs), but differ in their implementations (Kopylova, Navas-Molina et al. 2016). QIIME1 reportedly produced inflated numbers of OTUs with standard parameters (Edgar 2017). The successor of QIIME1 is QIIME2 (Bolyen, Rideout et al. 2019) that includes novel approaches to identify representative sequences. QIIME2 computes error-corrected amplicon sequence variants (ASV) for Illumina read sequences utilizing the software packages Divisive Amplicon Denoising Algorithm (DADA2) (Callahan, McMurdie et al. 2016) or Deblur (Amir, McDonald et al. 2017). ASVs are generally considered to be a more detailed view of OTUs as produced by QIIME1 or Mothur (Callahan, McMurdie et al. 2017). Ideally, ASVs represent actual amplicon sequences with single-nucleotide resolution that originate from each 16S rRNA gene copy of each species so that one species might be represented by several ASVs (Větrovský and Baldrian 2013). Still, 16S rRNA genes that do not differ in their amplified sequence cannot be resolved. MEGAN (MEtaGenome ANalyzer) was continuously maintained for more than a decade (Huson, Auch et al. 2007, Huson, Mitra et al. 2011, Huson, Beier et al. 2016). Initially, MEGAN was designed for metagenome analysis, but was also adapted to analyze 16S rRNA gene amplicon sequencing data (Mitra, Stärk et al. 2011). MALT (MEGAN ALignment Tool) is an extension of MEGAN and assigns sequencing data into taxonomic bins by sequence alignment (Herbig, Maixner et al. 2016). MALT with MEGAN does not compute representative sequences such as OTUs or ASVs but only taxonomic bins with abundances. Recovering representative sequences such as ASVs or OTUs allows for further analysis like constructing phylogenetic trees or performing targeted analyses, such as searching for the same sequence or related sequences in other data sets, that is not possible when using MEGAN. However, taxonomic analysis involving MEGAN can be straight forward with functional gene sequences that do not have elaborate reference databases like the 16S rRNA gene (DeSantis, Hugenholtz et al. 2006, Quast, Pruesse et al. 2013), e.g. for methane monooxygenase genes (*pmoA*, *mmoX*) or methanol or for dehydrogenase genes (*xoxF4*, *xoxF5*, *mxaF*) (Taubert, Grob et al. 2019). This type of functional gene analysis would require more efforts using QIIME or Mothur.

Various bioinformatics tools complicate the selection and the choice of the analysis approach. Open questions regarding the analysis of 16S rRNA amplicon sequences are: Which method performs best in respect to sequence recovery, relative quantification, and taxonomic classification and how do the obtained results differ between methods. In this study, we therefore aimed to i) identify the most suitable current bioinformatics method to examine environmental microbial communities based on 16S rRNA gene amplicon sequencing data with a focus on taxonomic identification and microbial diversity analysis and to ii) reveal differences caused by analysis pipelines. Key elements for our comparisons include the accuracy of recovered 16S rRNA gene amplicon sequences, their taxonomic classification and their quantification. All these elements are essential for exploring microbial communities, predicting ecological relevance, identifying biochemical key players or drawing conclusions about differences between communities. Here, we compared the most widely used 16S rRNA gene amplicon sequencing analysis tools with mock datasets and environmental samples and implement the findings as an nf-core workflow (Ewels, Peltzer et al. 2019, Straub and Peltzer 2019) to allow for execution in highly parallelized computing infrastructures, such as high-performance computing environments or compute clouds. Nf-core workflows strictly follow the FAIR (Findable, Accessible, Interoperable, and Re-usable) principle (Wilkinson, Dumontier et al. 2016), come with high quality standards, and are fully based on open source (Ewels, Peltzer et al. 2019).

## 2 Results

### Mock datasets showed highest sensitivity with QIIME1 but highest specificity with QIIME2

To evaluate the performance of the 16S rRNA gene sequencing analysis tools, three mock datasets (i.e. Balanced, Extreme, and HMP) based on samples with known composition were analyzed with Mothur, QIIME1, QIIME2, and MEGAN. First, the number of recovered 16S rRNA gene amplicon sequences (i.e. OTUs or ASVs) were compared to expected numbers, determined based on the reference sequences and abundances, and used as the basis for subsequent analyses. Only QIIME1, Mothur and QIIME2 generated sequences that could be compared to defined mock communities. MALT with MEGAN did not generate sequences and, therefore, did not allow this comparison. The number of OTUs or ASVs generally overestimated the number of expected unique 16S rRNA gene amplicons for all three datasets (Table 1). Mothur and QIIME1 in particular calculated 8- to 320-fold more sequences than expected, with 97% clustering similarity being at the lower end and 99% at the upper threshold. The number of sequences was much better estimated by QIIME2 in combination with DADA2 or with Deblur, however, Deblur underestimated the number of sequences for the Extreme dataset by almost 30% (Table 1). The accuracy of recovered 16S rRNA gene sequences and the relative sequence abundance is of particular interest for subsequent taxonomic classification or phylogenetic tree construction as well as for a realistic representation of microbial community composition. In the three mock datasets, QIIME1 showed highest sensitivity and recovered 83% to 94% of the reference sequences, closely followed by QIIME2 using DADA2 with 71% to 95% recovered sequences (Fig. 1, Table S1). The lowest sensitivity with only 43% (15 of 35 total; Table S1) recovered sequences was found for QIIME2 using Deblur while processing the Extreme dataset, where mainly low abundant sequences failed to be recovered (Fig. 1B).

**Figure 1:**
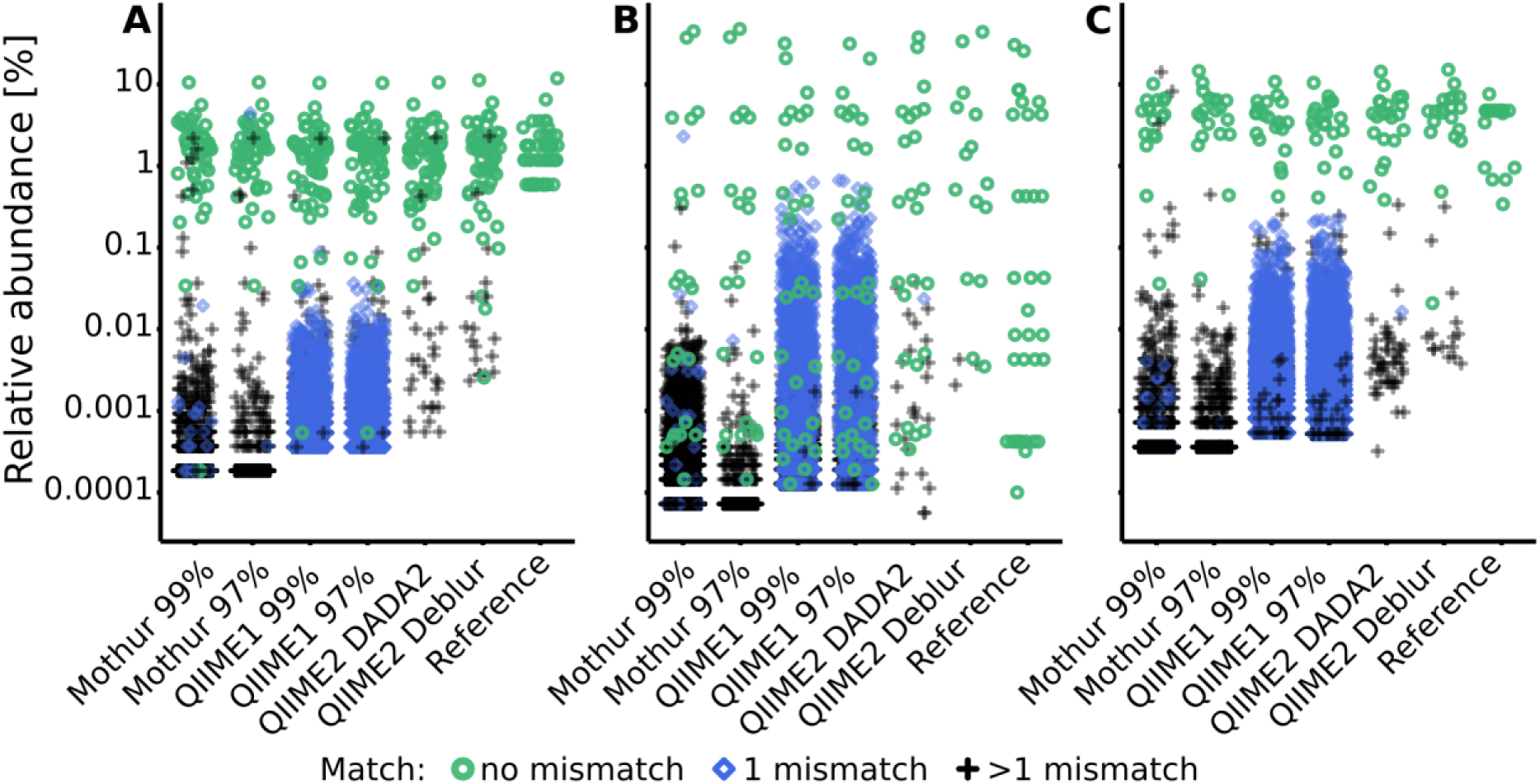
Relative abundance of sequences (i.e. OTUs or ASVs) and their distance to reference sequences for each pipeline and dataset (A: Balanced, B: Extreme, C: HMP). Sequences with perfect match to a reference sequence are depicted as green circles, with one mismatch as blue diamonds and with more than one mismatch as black crosses. Note that “1 mismatch” blue diamonds are covering most “>1 mismatch” black crosses for QIIME1. In the Jitter plot, noise (a random value) is added only to x-axis to separate dots.

**Table 1:** Generated sequences (i.e. OTUs or ASVs) for each analysis pipeline (Mothur, QIIME1, QIIME2) and mock dataset (Balanced, Extreme, HMP).

|  |  | Balanced | Extreme | HMP |
| --- | --- | --- | --- | --- |
| Species |  | 57<sup>a</sup> | 21 | 27 |
| Unique amplicons |  | 62 | 25 | 35 |
| Mothur | 99<sup>b</sup> | 1681 | 8009 | 1556 |
|  | 97<sup>b</sup> | 527 | 367 | 546 |
| QIIME1 | 99<sup>b</sup> | 1474 | 5545 | 4161 |
|  | 97<sup>b</sup> | 1002 | 1411 | 2826 |
| QIIME2 | DADA2<sup>c</sup> | 89 | 50 | 73 |
|  | Deblur<sup>c</sup> | 74 | 18 | 35 |

<sup>a</sup>57 species but only 52 distinguishable with the sequenced amplicon.
<sup>b</sup>similarity (%) at which sequences are clustered into operational taxonomic units (OTUs)
<sup>c</sup>ASV calling software

QIIME2 in combination with Deblur was most specific for all datasets with only 3 to 18 unexpected sequences, followed by QIIME2 in combination with DADA2 that produced 25 to 50 sequences that did not perfectly match to a reference sequence. However, of the 30 unexpected sequences found by QIIME2 with DADA2 in the Balanced dataset (Fig. 1, Table S1), 13 were reported by all pipelines, 10 were detected by all but one pipeline, and only 3 sequences were found exclusively by QIIME2 with DADA2 and no other pipeline. QIIME1 and Mothur detected at least 10 times more unexpected sequences than QIIME2 (Fig. 1, Table S1). For all investigated pipelines (excluding MEGAN), unexpected sequences mostly occurred below 1% relative sequence abundance, but the majority of unexpected sequences occurred below 0.001% to 0.1% abundance, depending on the pipeline (with QIIME1 at the higher end) and the dataset. For example, the Balanced dataset analyzed with QIIME1 had the majority of non-perfect matching sequences present at less than 0.01% relative abundance but the Extreme dataset at less than 0.1% relative abundance (Fig. 1). Using a 99% similarity threshold for OTUs with Mothur or QIIME1 did not improve sensitivity compared to the 97% similarity threshold but was highly detrimental to specificity by increasing the number of unexpected sequences by 50% to 2,000% (Fig. 1, Table S1).

### Taxonomic representation of the mock datasets was best resembled by QIIME2

To assess the accuracy of taxonomic classification, the F-score (Kopylova, Navas-Molina et al. 2016), that is the harmonic mean of precision (detected reference taxa to all *predicted* taxa) and recall (detected reference taxa to all *reference* taxa), was calculated for several taxonomic levels (i.e. class, order, family, genus and species). Taxonomic classification varied substantially among pipelines, for instance using the HMP dataset F-scores from 0.2 (Mothur) to 0.8 (QIIME2 in combination with Deblur) were generated at genus level (Fig. 2). Generally, QIIME2 had either close to the highest or the highest F-score of all four analysis pipelines in all datasets (Fig. 2), meaning that the compromise between precision and recall was best for QIIME2. Among all investigated pipelines, F-scores were similar for the Balanced dataset, but QIIME1 and QIIME2 achieved best results (i.e. highest F-score) for the Extreme datasets and QIIME2 for the HMP dataset above species level, i.e. genus level and higher. This difference was mainly driven by the superior precision of QIIME2 that was determined for all investigated datasets and for all taxonomic levels above species level. QIIME2’s Deblur outperformed DADA2 slightly on the Balanced dataset and more pronounced with the HMP dataset but had a lower F-score on family and genus level of the Extreme dataset due to Deblur’s higher precision but lower recall.

**Figure 2:**
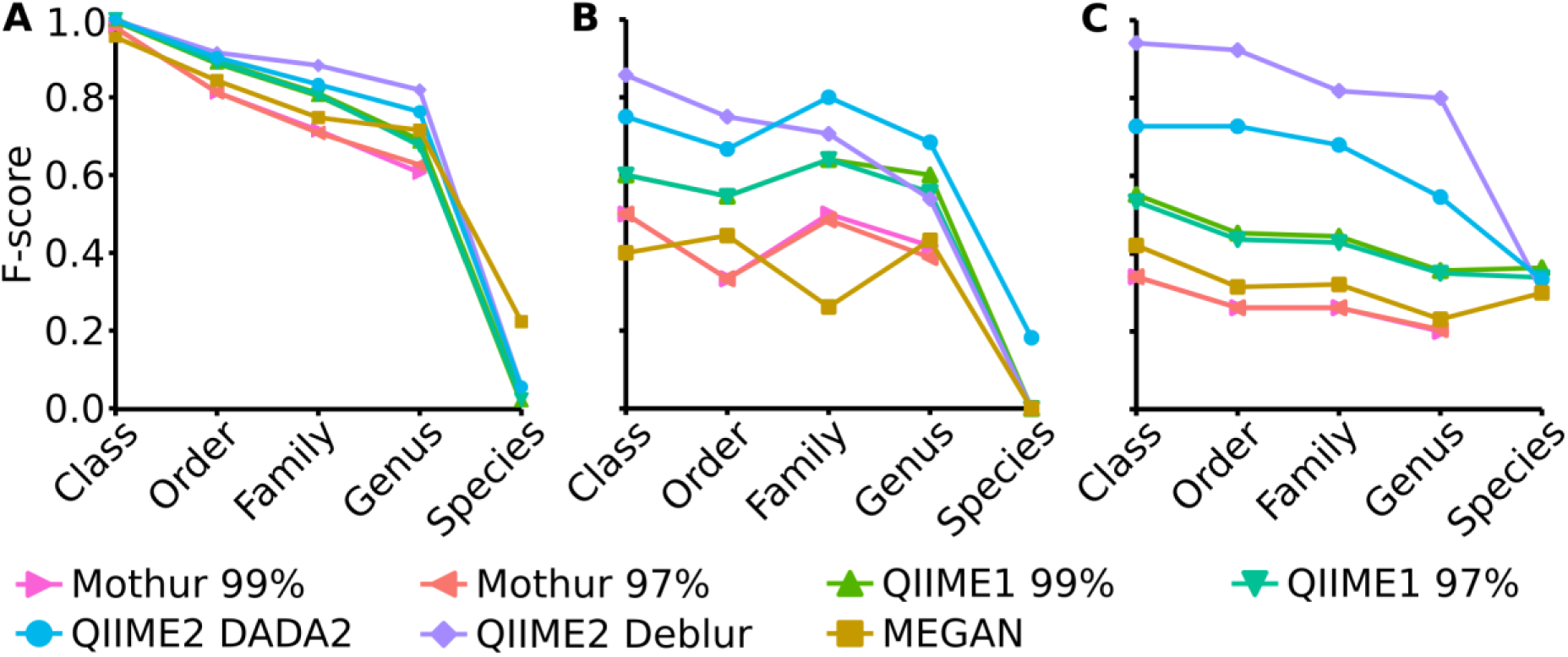
F-score on several taxonomic levels (i.e. class, order, family, genus and species) for Balanced (A), Extreme (B) and HMP (C) mock datasets.

Mothur and MEGAN achieved the lowest F-score for all taxonomic levels. In order to optimize the taxonomic classification with MEGAN, its 16S Percent Identity Filter was enabled and taxonomic assignments were projected to respective ranks, however, this did not improve taxonomic classification substantially compared to default settings. In fact, until genus level, the F-score for all three mock datasets were almost identical to those calculated with default settings but the species classifications were improved to the best values among all pipelines for Balanced and Extreme datasets, however, worsened for HMP data.

Generally, F-scores at species level were very low compared to higher taxonomic ranks. Mothur did not attempt to annotate the species rank at all unlike QIIME1, QIIME2 and MEGAN. Overall, the HMP dataset had the highest species annotation score of the three datasets with very similar values for all four analysis pipelines that annotated species (Fig. 2).

### Alpha-diversity indices of the mock datasets were approximated most closely by QIIME2

The Shannon index (Shannon 1948) that determines how many different types of species or sequences (i.e. OTUs or ASVs) are present in a sample (richness) and how evenly these are distributed (evenness) followed the expected trend for Mothur and QIIME2, i.e. the diversity decreased from Balanced (expected: 3.79) to HMP (expected: 3.09) to Extreme datasets (expected: 1.98). However, QIIME1 surprisingly led to a higher Shannon index for the HMP dataset (4.04) than for the Balanced dataset (3.90). The Shannon index of the Balanced dataset was relatively independent of the analysis pipeline and varied only slightly from 3.68 (Mothur 97%) to 3.93 (QIIME1 99%) when excluding the outlier MEGAN (3.06). But, for the HMP and Extreme datasets the pipelines came to different results with 2.42 (MEGAN) to 4.04 (QIIME1 99%) for the HMP dataset and 1.01 (MEGAN) to 2.98 (QIIME1 99%) for the Extreme dataset. Generally, QIIME1 overestimated the Shannon index for all mock datasets, while QIIME2 and Mothur slightly underestimated the values and MEGAN heavily underestimated the diversity in all datasets by 20% to 60% (Fig. 3). Enabling MEGAN’s 16S Percent Identity Filter shifted the calculated Shannon index closer to the expected values for Extreme data but further away for the other two datasets compared to default settings. However, the expected Shannon indices were calculated on sequence level and MEGAN used genera abundance estimates instead of fine grained OTU or ASV sequences and therefore was not able to closely resemble the expected numbers.

**Fig. 3:**
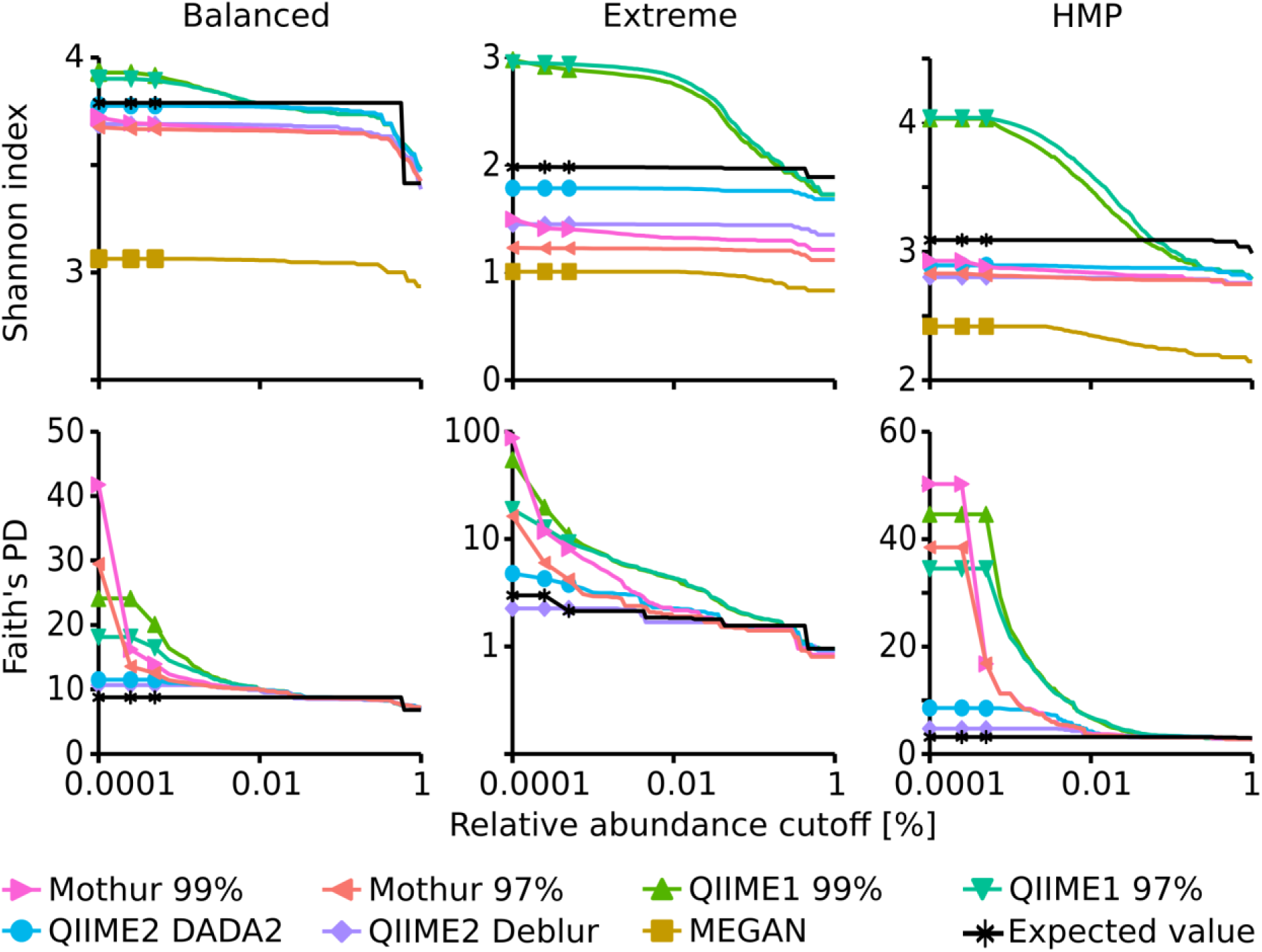
Alpha diversity indices dependent on a relative abundance cutoff (up to 1%). Top: Shannon index, Bottom: Faith’s Phylodiversity; note the logarithmic x-axis and for Faith’s PD of the Extreme dataset the logarithmic y-axis.

Overall, Shannon alpha diversity indices were most accurately reproduced by QIIME2 in combination with DADA2 though this method underestimated Shannon diversity on average by 6% (0.3% to 10% for the three datasets).

Faith’s Phylodiversity (PD) index (Faith 1992) that is a qualitative measure of the sum of the phylogenetic branch lengths covered by a sample was also best resembled by QIIME2 with almost 2-fold overestimation on average (1.3- to 2.7-fold for the three datasets) but strongly overestimated by QIIME1 (2- to 18-fold) and Mothur (3.5- to 29-fold). The estimates improved with increasing relative abundance cutoff and resembled the expected values very closely, ranging from 0.9- to 1.2-fold of the expected values when only sequences above 0.1% abundance were considered (Fig. 3).

### Poor agreement in recovery of 16S rRNA gene amplicon sequences and taxa of environmental samples between analysis methods

Environmental samples typically have a more complex microbial community than mock datasets and, therefore, we selected samples from diverse habitats (groundwater, soil, river sediment, and river water) for analysis using 16S rRNA gene sequencing to investigate whether results differed between the investigated pipelines. First, the numbers of reported sequences (i.e. OTUs and ASVs) and unique genera for each analysis pipeline were compared to investigate whether the trend observed in the mock datasets was also evident in the environmental samples, since measures such as estimates of community diversity or clustering distance strongly depend on sequence or taxa count. Total sequence numbers, e.g. OTUs or ASVs, across all samples varied from 11,747 with QIIME2 and Deblur to 79,326 with Mothur. This was a similar trend compared to the analysis of the mock datasets, where QIIME2 with Deblur and DADA2 produced the lowest amount of sequences (ASVs) while QIIME1 and Mothur counted the highest number of sequences (OTUs). Most sequences (10,223 ASVs) computed by Deblur (87%) or DADA2 (55%) were identical (Fig. 4A). However, there was relatively little overlap between Mothur, QIIME1, and QIIME2 with only 6,453 (6%) identical sequences (Fig. 4B), whereas 62,736 sequences (55%) were shared by at least two but not all pipelines and 44,702 sequences (39%) were not shared at all. 17,478 and 8,936 OTU sequences overlapped within Mothur or QIIME1 with varying similarity cutoffs (i.e. 97% or 99%), respectively, and both analysis pipelines produced 1.3- to 2-fold more OTU sequences with the 99% similarity cutoff than with 97%, in line with the findings of the mock community analysis that produced 1.5- to 23-fold more OTUs with 99% similarity cutoff.

**Figure 4:**
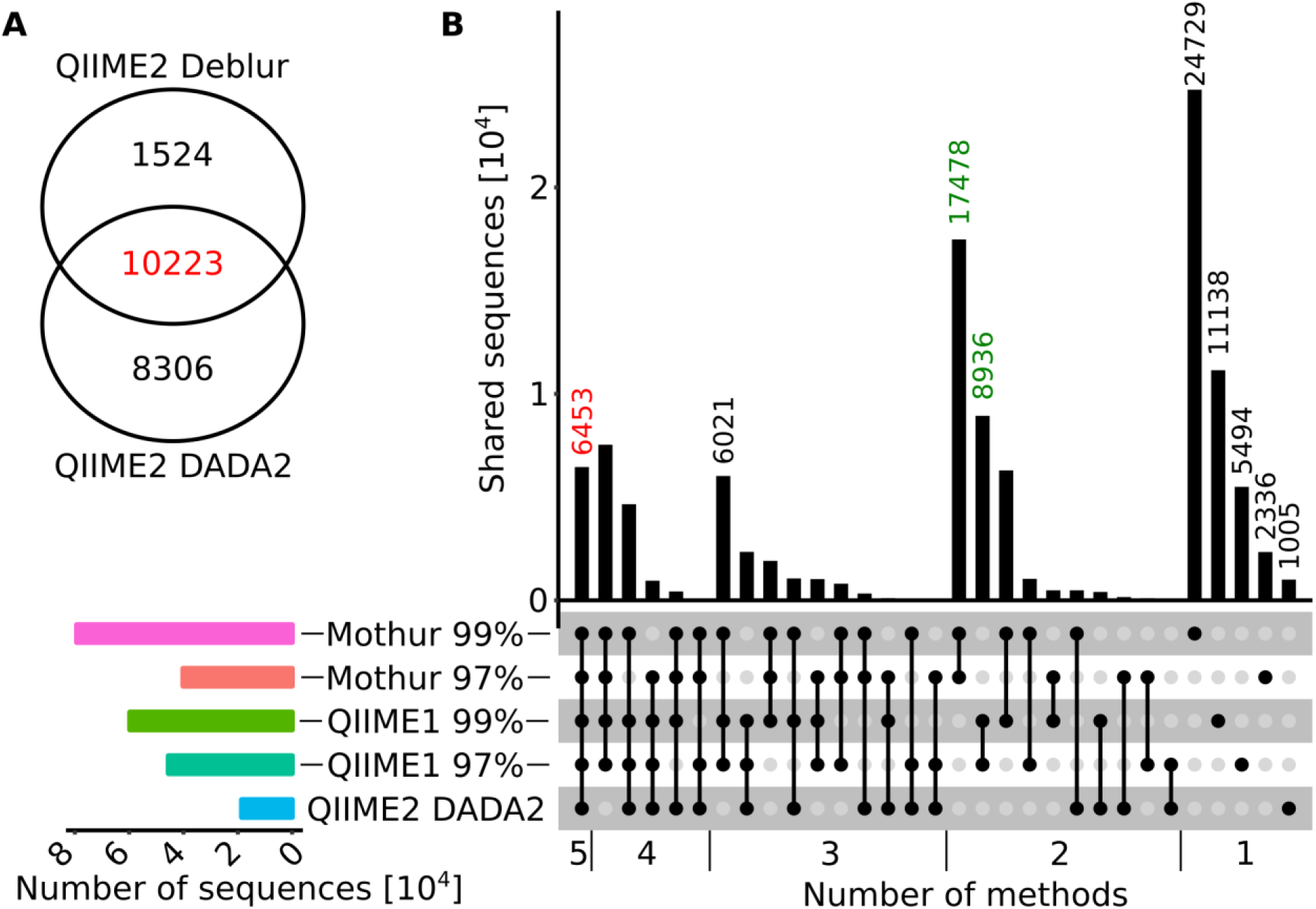
Overlap diagrams for ASV or OTU sequences. A: Venn diagram for QIIME2 with DADA2 and Deblur sequences that were trimmed to 250 bp for comparison. B: Horizontal bar plot (bottom, left panel) represents the number of sequences reported for each method (from top to bottom: Mothur 99%, Mothur 97%, QIIME1 99%, QIIME1 97%, QIIME 2 with DADA2) and matrix (bottom, right panel) indicates methods that are part of an intersection with (connected) black filled circles. The vertical bar graph (top, right panel) represents the number of shared sequences (intersection size) for each intersection. In the vertical bar graph, the number of sequences calculated by all methods are highlighted in red and sequences only found within QIIME1 or Mothur in green.

Reported genera derived from the up to 8-fold different sequence counts (11,747 to 79,326, Fig. 5A) varied much less, from 1,861 using QIIME2 with Deblur to 2,640 using QIIME1. MEGAN reported 961 genera, which was by far the lowest number (Fig. 5B). All pipelines recovered the highest number of sequences from sediment site 1 (26% to 44% of total) and least sequences in groundwater site 1 (QIIME1: 10% to 12% and QIIME2: 11%) or site 2 (Mothur: 9% to 13%). Most genera, however, were found in sediment site 2 (Mothur: 69% to 71%), river water site 1 (QIIME1: 64% to 66% and MEGAN: 61%) or sediment site 2 (QIIME2: 42% to 53%). But generally, all pipelines identified more sequences and genera in sediment and river samples than in groundwater and soil samples (Fig. 5).

**Figure 5:**
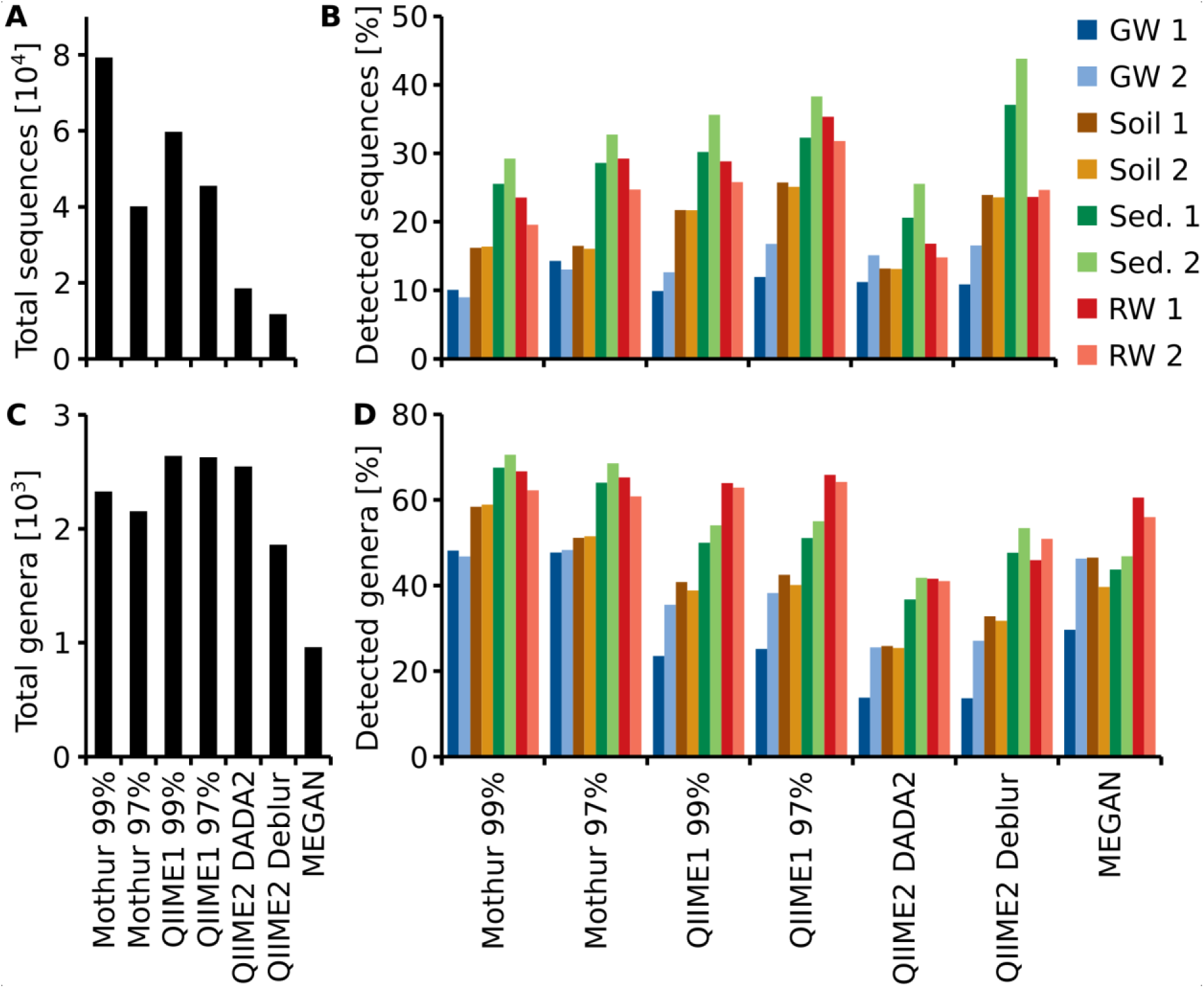
Total number of recovered sequences (OTUs or ASVs) (A) and genera (C), for each pipeline and percentage of total sequences (B) and genera (D) detected at each sampling site. For each habitat (groundwater, soil, sediment or river water), two sampling sites (1, 2) were investigated.

QIIME2 with DADA2 found on average the same number of ASVs per sampling site as QIIME2 with Deblur. However, QIIME2 with DADA2 found almost twice the number of ASVs overall compared to QIIME2 with Deblur (Fig. 5A). Thus, more unique sequences were found per sampling site with DADA2 while Deblur found the same ASVs at multiple sampling sites (Fig. 5A-B). The same was also true at genus level (Fig. 5C-D).

To see how the analysis methods differed in their taxonomic classification, the relative abundance at phylum and genus level was compared. Generally, at phylum level the community composition at each individual site was very similar regardless of the pipeline with one exception; at groundwater site 1, the community composition showed large differences between all tested pipelines at phylum level (Fig. 6A, Fig. S1). At genus level, however, dramatic differences in the resulting community composition were observed for all pipelines. For example, while the community composition at phylum level at soil site 1 was fairly consistent between all pipelines, the results at genus level were substantially different to the extent that the most abundant genera were not detected across all pipelines (Fig. 6B, Fig. S2). Some similarities were observed between QIIME pipelines, for example, the community composition at genus level in groundwater site 1 differed only slightly when using QIIME1 with either the 99% or 97% similarity threshold and QIIME2 with either Deblur or DADA2. In contrast, the differences between the QIIME pipelines and MEGAN varied substantially.

**Figure 6:**
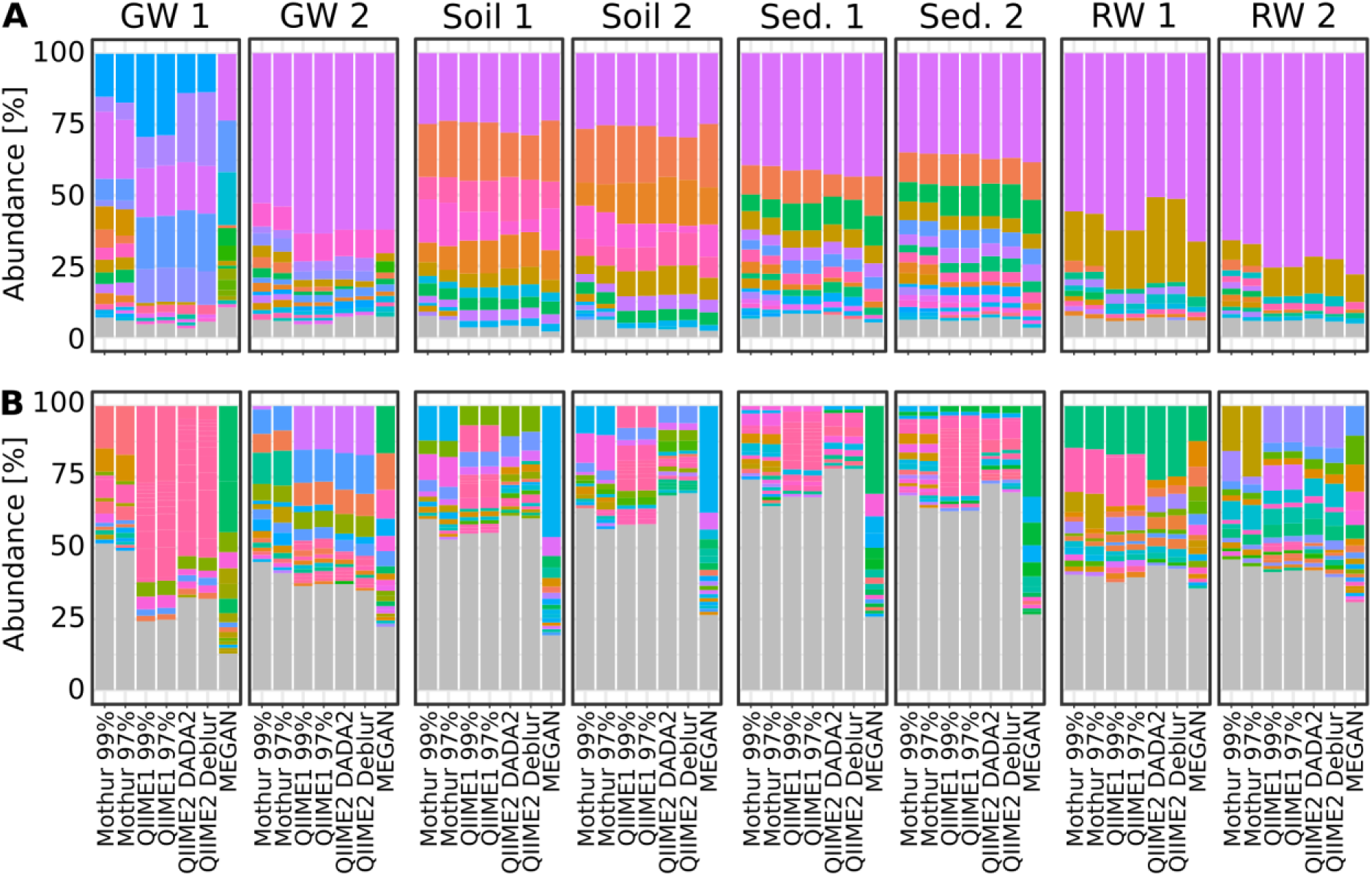
Bar plots showing relative taxa abundance averaged over triplicates at (A) phylum or (B) genus level for all habitat sampling sites and analysis methods. Genera or phyla are shown in the same colors throughout the figure. Grey are taxa <1% abundance. GW, groundwater; Sed., sediment; RW, river water; 1 & 2 indicate sampling sites.

Because dominant microbial taxa might have important functions at specific sites, the five most abundant taxa at genus level were compared for each sampling site and analysis method. QIIME1 with 99% or 97% similarity threshold identified always the exact same five most abundant taxa. QIIME2 using DADA2 agreed almost perfectly with QIIME2 using Deblur, differing only in two sampling sites by the fifth most abundant genus (soil site 2: DADA2: *Candidatus Nitrocosmicus* vs. Deblur: an uncultured *Burkholderiaceae*; river water site 2: DADA2: *Acinetobacter* vs. Deblur: *Arcobacter*; Table S2). Very little difference between the similarity thresholds of 99% and 97% was observed for Mothur which had on average more than four identical taxa in the top five most abundant ones across sampling sites (Fig. 7, Table S2). MEGAN had lowest agreement with all other methods and none or maximum two reported genera per sampling site matched those found using the three other methods (Fig. 7, Table S2). Only up to two overlapping taxa on average were observed by Mothur compared to QIIME1 as well as Mothur compared to QIIME2, and on average 2.8 overlapping taxa were found by QIIME1 compared to QIIME2. Differences were also observed in the consistency of the five most abundant genera between analyses depending on sampling site. For example, at river water site 1, the average number of overlapping most abundant genera was 3.0. Conversely, sediment site 1 had the lowest average number of overlapping genera with 0.9, showing a clear difference in the community composition results, depending on the analysis method (Table S2). However, across all samples, even when the same genera were found, the abundance and the order of abundance varied with the analysis method. For example, river water site 1 analyses showed that (except for MEGAN) *Rhodoferax*, *Malikia* and *Flavobacterium* were consistently present in the top five most abundant genera. However, of these three taxa, according to Mothur with 97% similarity threshold, *Malikia* had highest abundance and *Rhodoferax* the lowest, while according to all other methods *Rhodoferax* had the highest abundance and *Flavobacterium* the lowest. Additionally, the relative abundance for *Malikia* varied from 5% (QIIME1) to 9% (Mothur). On the other hand, most analyses agreed about the order of abundance of the three most abundant taxa in groundwater site 2 which was dominated by *Gallionellaceae* (Mothur) or *Sideroxydans* (QIIME1 and 2), belonging to the family *Gallionellaceae*, followed by *Polaromonas* and *Acinetobacter*. In contrast to all other analyses, MEGAN reported an unclassified taxon, *Acinetobacter*, and *Candidatus Omnitrophica* as the three most abundant taxa in groundwater site 2 (Table S2).

**Figure 7:**
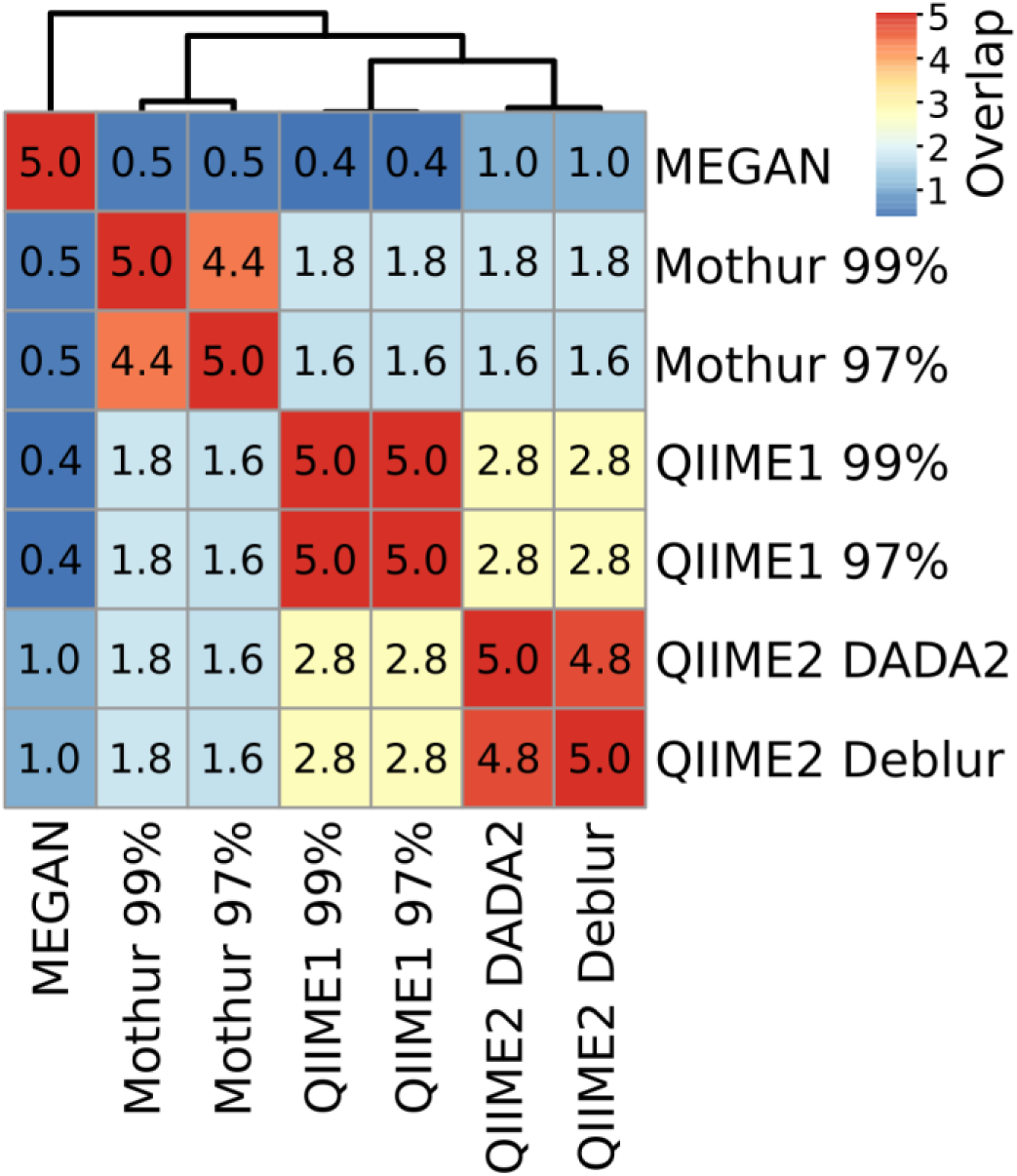
Heatmap and number of shared taxa among the top five most abundant genera for each pipeline averaged across sampling sites. Genus names and their relative abundance can be found in Table S2.

### Diversity estimates in environmental samples varied among tools

To investigate the comparability of within-sample diversity measures (alpha-diversity), the Shannon index was calculated for results of each analysis pipeline. Generally, across all samples, QIIME2 with DADA2 and with Deblur reported similar values (± 1%) to Mothur with 97% similarity threshold for the Shannon index. In comparison, QIIME1 and Mothur with 99% similarity threshold had 13% and 9% higher values, respectively, while MEGAN calculated 20% lower values (Fig. 8A). The trend seemed very similar among all analysis pipelines except for MEGAN (Fig. 8B-E), with descending diversity from sediment, to soil, to groundwater, and river water, although there were differences in absolute Shannon diversities. In contrast, MEGAN reported river water having the highest Shannon diversity followed by all other habitats (Fig. 8E).

**Figure 8:**
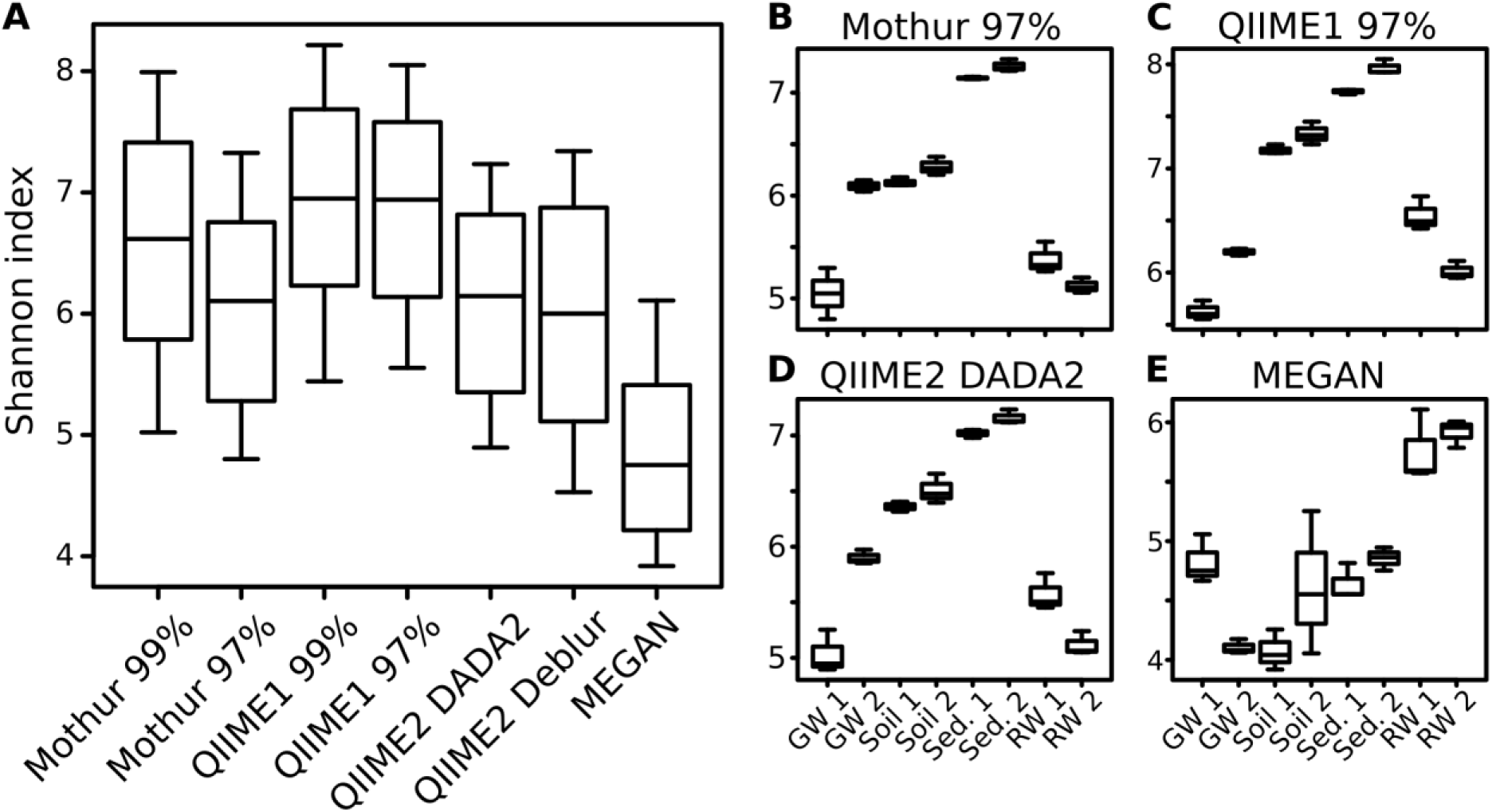
Comparison of alpha-diversity indices for environmental samples. For each analysis pipeline, Shannon indices were calculated either on all samples combined (A) or each sampling site separately (B, C, D, E): GW, groundwater; Sed., sediment; RW, river water; 1 & 2 indicate sampling sites.

To investigate if each pipeline allowed similar sample groupings, distances based on OTUs (Mothur and QIIME1), ASVs (QIIME2) or taxa (MEGAN) abundance between samples (beta-diversity) were measured. Overall, distances and groupings were similar across pipelines and all of them allowed the separation of habitats except MEGAN, where it was not possible to distinguish between samples from river water and groundwater as clearly as for other pipelines (Fig. 9A, arrow 1). Also, the two groundwater sites clustered separately by all pipelines (Fig. 9).

**Figure 9:**
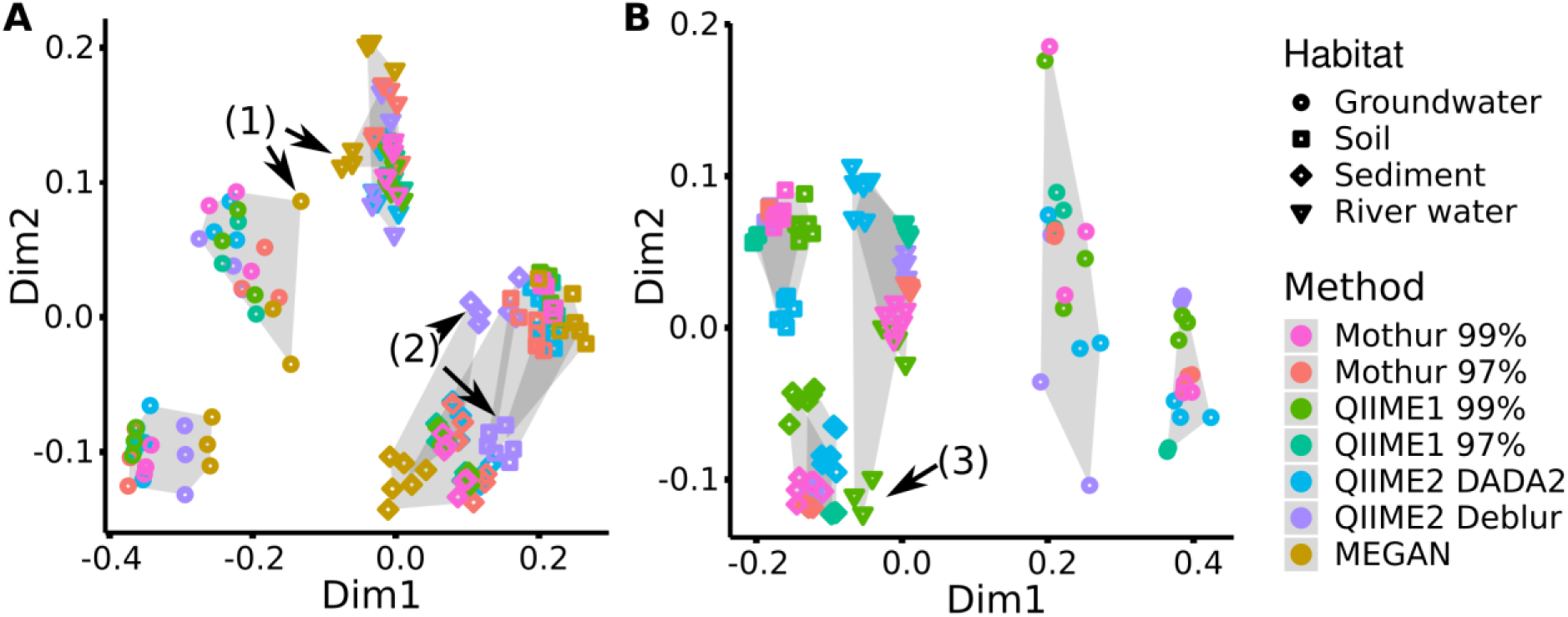
Comparison of beta-diversity plots for environmental samples. For each analysis pipeline, A: Bray-Curtis distance (quantitative) or B: Unweighted UniFrac distances (qualitative and phylogenetic) were subjected to NMDS (Non-metric Multidimensional Scaling) ordination and combined by Generalized Procrustes Analysis (NMDS stress values < 0.08, Table S3). Grey shading marks sampling sites, e.g. all three replicates analyzed by the different pipelines from one sampling site are connected by grey background.

In terms of consistency between pipelines, QIIME2 with Deblur exchanged placement of sediment and soil samples on the Bray-Curtis (Bray and Curtis 1957) distance plot compared to other pipelines but cluster separation of these two habitats remained stable (Fig. 9A, arrow 2). Calculation of Unweighted UniFrac (Lozupone, Hamady et al. 2007) requires sequences (i.e. ASVs or OTUs) so that MEGAN was excluded from the following comparison: QIIME1 with 99% similarity threshold placed the samples from a river site and the sediment samples differently (Fig. 9B, arrow 3), while QIIME2 in combination with DADA2 had slightly shifted placement for river and soil samples, resulting in a slightly rearranged plot for both methods compared to all other pipelines (Fig. 9B).

## 3 Discussion

Here we compare 16S rRNA gene sequence analysis pipelines and aim at identifying the best suited bioinformatics method to date to analyze environmental microbial communities based on high-throughput DNA- or RNA-based 16S rRNA (gene) amplicon sequencing data. Therefore, mock communities and environmental samples from a range of habitats with differing geochemical conditions (e.g. redox and nitrate concentrations) were analyzed with popular analysis pipelines, i.e. Mothur, QIIME (version 1 and 2) and MEGAN.

### The best compromise of sensitivity and specificity by QIIME2

Generally, Mothur and QIIME1 recovered almost all 16S rRNA gene amplicon sequences and genera but the number and abundance of false positives was relatively high, so that sometimes the true positive sequences were buried in a huge number of false positives. Removing sequences with low relative abundance, e.g. <0.1%, improved the results for QIIME1 and Mothur but had the adverse effect of removing low abundant expected sequences. Low abundant sequences and taxa might be interesting in some studies, e.g. when a group of low abundant taxa is performing a crucial step in the biogeochemical cycle, such as carbon and nitrogen cycling or sulfate reduction (Musat, Halm et al. 2008, Pester, Bittner et al. 2010, Jousset, Bienhold et al. 2017), and therefore removing them might be undesirable. QIIME2 using Deblur suffered from relatively low recall (several sequences or taxa were not detected) but had highest precision (a low number of additional false sequences or taxa was detected) that was similar to findings observed by Nearing *et al.* (2018). A recent study found that DADA2 had difficulties finding low abundant variants and produced few but high abundant false positives (Hathaway, Parobek et al. 2017), although we did not observe this finding. On the contrary, QIIME2 using DADA2 showed high recall and high precision. However, in the Balanced mock dataset QIIME2 found highly abundant unexpected sequences, but because these were in the majority (23 of 30 total) also found by all or all but one other method, we assumed these were true sequences not present in the reference database. Essentially, perfect results were not obtained by any method but DADA2 in combination with QIIME2 seemed the best compromise of sensitivity and specificity (Table 2).

**Table 2:** Summary of strengths and weaknesses of tested pipelines. All values are means (± standard deviation) of the analyses of three mock datasets.

|  | Mothur |  | QIIME1 |  | QIIME2 |  | MEGAN |
| --- | --- | --- | --- | --- | --- | --- | --- |
|  | 99<sup>a</sup> | 97<sup>a</sup> | 99<sup>a</sup> | 97<sup>a</sup> | DADA2<sup>b</sup> | Deblur<sup>b</sup> |  |
| Precision (%) | 1 ± 1 | 6 ± 3 | 2 ± 2 | 3 ± 3 | 49 ± 18 | 72 ± 13 | n.d. |
| Sensitivity (%) | 69 ± 7 | 72 ± 11 | 87 ± 6 | 87 ± 6 | 85 ± 12 | 71 ± 25 | n.d. |
| Taxonomy<sup>c</sup> | 49 ± 23 | 49 ± 22 | 63 ± 18 | 63 ± 19 | 77 ± 8 | 80 ± 9 | 44 ± 27 |
| Shannon<br>index<sup>d</sup> | -11 ± 12 | -16 ± 19 | 28 ± 23 | 28 ± 23 | -6 ± 5 | -13 ± 13 | -30 ± 17 |

n.d.: not determined by this method.
<sup>a</sup>similarity (%) at which sequences were clustered into operational taxonomic units (OTUs)
<sup>b</sup>ASV calling software
<sup>c</sup>F-score on family level in percent
<sup>d</sup>in % deviation; negative numbers mean underestimation, positive numbers mean overestimation

Taxonomic annotation depends on the amplicon region (Kozich, Westcott et al. 2013), reference database, and the classifier (Almeida, Mitchell et al. 2018). The reference database used in this study was SILVA v132 with 16S rRNA gene sequences dereplicated at 99% similarity, meaning it contained combined taxa with ≥99% similar 16S rRNA gene sequences and thereby reduced the computational requirements. However, it also decreased taxonomic resolution. The mock datasets used here contained sequences of the 16S rRNA gene V4 region with a length of 250 to 254 bp. The choice of the amplified region also restricts taxonomic resolution, e.g. the *Enterobacteriaceae* family and the *Clostridiales* order are known to be poorly resolved using these short V4 amplicons (Jovel, Patterson et al. 2016) and the resolution at phylum level is lower than sequencing the whole 16S rRNA gene (Yang, Wang et al. 2016). But even when using full-length 16S rRNA gene analysis, some related but distinct microorganisms can remain unresolved. For instance, five *Streptomyces* species with identical 16S rRNA gene sequences were shown to have phenotypic, microscopic, genetic and genomic differences (Antony-Babu, Stien et al. 2017). Overall, our study showed that species level seemed too biased to be trusted for taxonomic classification. This is in agreement with earlier studies that found species classification unreliable especially for uncharacterized species (Bokulich, Kaehler et al. 2018, Edgar 2018) but taxonomic classification at genus level was more accurate.

### The choice of the analysis pipeline affects the outcomes of studies

The difference in the number and the quality of recovered 16S rRNA gene amplicon sequences and their further taxonomic classification among pipelines also caused deviations in data interpretation. For instance, the sampling site with the highest microbial diversity among the investigated environmental samples (i.e. groundwater, soil, sediment or river water sites) differed depending on the analysis pipeline. Additionally, differences in microbial diversity estimates led to dissimilar interpretations depending on the analysis pipeline. The choice of the analysis pipeline affected the outcome of our study and, thus, special care needs to be taken when interpreting results, particularly when dealing with highly diverse environments.

#### Sequence recovery

Overall, the accuracy of sequence recovery of QIIME2 indicated that this pipeline was the best basis for further downstream analysis and data interpretation. This was due to denoising (i.e. DADA2, Deblur) that performed better than OTU clustering (i.e. Mothur, QIIME1), in line with other studies (Callahan, McMurdie et al. 2017, Nearing, Douglas et al. 2018). In contrast to Deblur, which uses a static error model to correct raw sequences, DADA2 computes an error model for each sequencing run based on potentially all samples (up to 1 million reads), requiring a re-analysis when only a subset of the initial samples is used in the final reporting. As a consequence, DADA2 requires much more computing time. However, Deblur will miss all amplicons that fall below a required length truncation threshold (e.g. 250 bp in this study) because all shorter amplicons are discarded, thus 1.36% of all sequences in the SILVA v132 database are ignored (Fig. S3).

Furthermore, all amplicons that are longer than the length threshold of 250 bp are cut and therefore essential data is lost. For example, 70% of all sequences in SILVA v132 (99% identity clustered and V4 region extracted) are 253 bp long and are therefore cut by 3 bp, losing >1% of data (Fig. S3). On the other hand, DADA2 requires choosing read-trimming cutoffs according to data quality, however, there are no defined rules for selecting these cutoffs and, without having a clear expectation of the result, it appears impossible to find the optimal solution. Essentially, operating Deblur seemed riskier than DADA2 because sequences that are below a chosen cutoff can be lost and overlooked using Deblur. Another advantage of DADA2 in our study was the high proportion of recovered sequences and taxa that were specific for each environmental sampling site. Nearing *et al.* (2018) observed the same trend and suspected that this was due to DADA2’s unique way to create pooled error profiles followed by sample-by-sample ASV picking. This implies that DADA2 might be better in separating similar sequences from different samples than tools that pick sequences from pooled samples (e.g. Deblur, QIIME1, Mothur), however, it is not possible to test this hypothesis with the investigated datasets in the present study.

#### Taxonomic identification

At genus level, there were substantial differences in the taxonomic overview (presented as bar plots), particularly for the top five most abundant genera, that each method provided. While mock datasets are often analyzed at lower phylogenetic levels, e.g. genus (Almeida, Mitchell et al. 2018, Nearing, Douglas et al. 2018), environmental datasets are also often shown at higher levels, e.g. phylum (de Voogd, Cleary et al. 2015, Oliveira, Gunderman et al. 2017). This might be due to the increasing complexity of graphs with increasing microbial diversity. For example, genera below one percent abundance accounted for 75% of the total abundance in the highly diverse soil and sediment habitats, investigated in this study, and were better represented by higher taxonomic levels such as phylum, where less than 10% abundance was accounted for by taxa with less than 1% abundance. However, in lower diversity habitats, i.e. groundwater and river water, the majority of genera were present at above one percent abundance and were reasonably well represented in stacked bar graphs at genus level. Of great concern is the low reproducibility among methods at genus level compared to phylum level. Showing low taxonomy levels down to genus (but not species) was only acceptable when using denoisers, i.e. QIIME2 with DADA2 or Deblur, and should be approached with caution when using OTU picking methods, i.e. Mothur and QIIME1, or taxonomic binning by MEGAN. This is because OTU methods and taxonomic binning performed worse on mock datasets than denoisers, and denoisers reported very similar genera for individual environmental samples. The accuracy of the taxonomic representation decreased with decreasing taxonomic ranks and was best for QIIME2 until genus level but was unreliable at species level for all methods. Better taxonomic resolution and classification might be achieved by investigating a larger fraction of the genome such as the full 16S rRNA gene. The V4 sub-region is a good choice because it allows complete overlap of paired-end sequences, thus reducing sequencing errors (Kozich, Westcott et al. 2013), and it closely resembles the phylogenetic signal of the whole 16S rRNA gene (Yang, Wang et al. 2016). The V4 sub-region was therefore also the focus of this study. Compared to sequencing a short region of the 16S rRNA gene with Illumina technology, whole 16S rRNA gene sequencing with Pacific Biosciences (PacBio) technology generates better results in terms of taxonomic resolution (Schloss, Jenior et al. 2016). PacBio circular consensus sequences (CCS) are produced by reading a circular short sequence (1 to 20 kb), such as the full 16S rRNA gene sequence, several times, thus achieving comparably low error rates similar to Illumina sequencing (Singer, Bushnell et al. 2016). High quality analysis is promised through DADA2 that was recently adapted to be able to denoise PacBio CSS (Callahan, Wong et al. 2019). However, PacBio CSS technology is currently not competitive in terms of sequencing depth, price, or availability. Targeting the even longer 16S-ITS-23S sequences of the *rrn* operon with Oxford Nanopore Technologies (ONT) sequencing allowed a high resolution at species level in a recent study (Cuscó, Catozzi et al. 2018). ONT sequencing is continuously improving and, similar to PacBio’s CSS technology, consensus reads are enhancing the accuracy of amplicon sequencing by a large margin, however, ONT’s sequencing accuracy is currently still considered inferior compared to Illumina or PacBio (Calus, Ijaz et al. 2018).

#### Alpha-diversity

The Shannon index is relatively insensitive to low abundant features (i.e. OTUs, ASVs or taxa) because it uses quantitative information (Shannon 1948) and the best possible estimates of Shannon diversity indices were calculated based on QIIME2 using DADA2. The closest resemblance of Faith’s PD required filtering for above 0.01% relative abundance for QIIME2 using Deblur or above 0.1% for QIIME2 using DADA2. Faith’s PD is a qualitative measure (Faith 1992) and therefore sensitive to the number of features irrespective of their abundance. Qualitative measures are better estimated on high confidence (e.g. high abundant) features, especially for error-prone OTU methods (Bokulich, Subramanian et al. 2013). Taking into account the high number of low-abundance, false-positive sequences in our study, quantitative diversity indices should always perform better on unfiltered data than qualitative measures. An unsuitable approach was to simply count OTUs/ASVs as diversity estimator because this resulted in an overestimation (QIIME1, Mothur, QIIME2 using DADA2) or in an underestimation for low abundant expected taxa (QIIME2 using Deblur).

#### Beta-diversity

Similar to alpha diversity measures, quantitative beta diversity methods are expected to perform better than qualitative ones, when expecting inaccurate, low abundant features (Lozupone, Hamady et al. 2007). However, in this study quantitative Bray-Curtis distances showed a similar sample discrimination as the qualitative Unweighted UniFrac distances. Unweighted Unifrac ignores relative abundances but takes phylogenetic distances into account and, thus, interprets phylogenetically similar sequences between samples as a smaller beta-diversity distance compared to phylogenetically distant sequences.

Beta-diversity distances were relatively similar between analysis methods despite the high variability in taxonomic classification. The underlying data structure (i.e. raw sequencing reads, OTU or ASV) for calculating beta-diversity distances is generally similar but mapping sequences to taxonomies performed differently. This is because the methods use very different approaches to resolve taxonomic classification (Almeida, Mitchell et al. 2018). These differences in taxonomic classification are expected to be larger for complex communities with sequences that are poorly represented in databases such as environmental samples and smaller for well-characterized communities such as those stemming from the human gut.

## Conclusion

Overall, we found that the results of Mothur, QIIME1, QIIME2, and MEGAN varied in all aspects but were generally more similar for high abundance features (i.e. OTUs, ASVs or taxa) and at higher taxonomic levels (e.g. phylum) and less similar for lower (<0.1%) abundant features and at lower taxonomic levels (e.g. genus). Therefore, biological indicators such as richness estimators or diversity indices differed between analyses, skewing conclusions. Yet, the “rare biosphere” is of particular interest for many studies (Musat, Halm et al. 2008, Pester, Bittner et al. 2010, Jousset, Bienhold et al. 2017) and taxonomic classification is often desired beyond genus level, making the selection of the best pipeline a crucial step. Based on our comparison, none of the investigated analysis methods was optimal, but QIIME2 performed best though it was not clear whether Deblur or DADA2 performed better in combination with QIIME2. However, the loss of data and amplicon sequences, that were below a certain length threshold, with Deblur seemed more daunting than choosing DADA2’s sequencing read truncation cutoffs. Because the longer runtime of DADA2 is of low importance when using high-performance computer clusters as well as because of the high sensitivity of DADA2, which appeared a better choice than the lower sensitivity but slightly higher specificity of Deblur, QIIME2 using DADA2 appeared as the best compromise (Table 2). The lessons learned in this study were implemented in the 16S rRNA gene amplicon sequence analysis pipeline found in the nf-core collection (https://nf-co.re) (Ewels, Peltzer et al. 2019), named nf-core/ampliseq (Straub and Peltzer 2019), to support data analysis that follows FAIR principles (Wilkinson, Dumontier et al. 2016). The pipeline has extensive documentation, requires minimal user input and no software installation except Java (which is available on most systems anyway), Nextflow (workflow management) and Singularity (software container management). It is independently citable using a Zenodo DOI (Straub and Peltzer 2019). All required software dependencies are bundled in containers and are automatically used by this workflow whenever an analysis is performed with a pipeline release. This ensures a fully reproducible analysis of high-throughput DNA- or RNA-based 16S rRNA (gene) amplicon sequencing data for future studies.

## 4 Methods

### Mock test datasets

To compare the performance of Mothur, QIIME1, QIIME2 and MALT/MEGAN, three mock datasets differing in microbial community composition, abundance distribution, and data quality were selected. All three datasets investigated the V4 region of the 16S rRNA gene and were sequenced by Illumina MiSeq. The “Balanced” dataset consisted of 57 bacteria and archaea from a broad range of habitats at even amounts of purified genomic DNA (Schirmer, Ijaz et al. 2015). The “Extreme” community contained 27 human gastrointestinal tract bacterial isolates at frequencies spanning six orders of magnitude and differing by as little as one nucleotide, for which 16S rRNA gene amplicons of individual cultures were quantified and pooled (Callahan, McMurdie et al. 2016). The “HMP” (Human Microbiome Project) dataset contained 21 well-separated strains in the human body with equimolar concentrations of 16S rRNA gene copies per genome (Kozich, Westcott et al. 2013); the sequence quality was the lowest of the three mock datasets. The Balanced dataset was retrieved from the European Nucleotide Archive (ENA), study PRJEB6244 sample M35 (SAMEA3298272) (Schirmer, Ijaz et al. 2015), the Extreme dataset SRR2990088 (Callahan, McMurdie et al. 2016) was downloaded from the Sequence Read Archive (SRA), and the HMP dataset, alias Mock1, was downloaded at mothur.org in set “130403” (Kozich, Westcott et al. 2013). For the Balanced and HMP datasets, primers were removed from raw sequences and untrimmed sequences were discarded with Cutadapt v1.14 (Martin 2011) wrapped by Trim Galore! v0.4.5 (Felix Krueger, bioinformatics.babraham.ac.uk).

For the Extreme and HMP datasets, reference sequences were obtained from the supplementary file “Supplementary Software” from Callahan *et al.* (2016). For the Balanced dataset, reference sequences for each species were retrieved from the “ribosomal RNA operon copy number database” (rrnDB) v5.4 (Stoddard, Smith et al. 2015). For QIIME2 in combination with Deblur, all recovered amplicon sequences had equal lengths determined by an input parameter and, therefore, reference sequences were truncated to the same length.

Further details on mock datasets and trimmed primers can be found in the supplementary information (Table S4).

### Sample Collection and DNA Extraction

Groundwater, soil, river sediment and river water were each sampled in triplicate at two sites.

#### Groundwater

Groundwater was collected from the monitoring well in Haslach (sampling site 1) using a submersible pump operating at the top of the screened section in 70 m depth below the well head (47.7 m below the water table) with a flow rate of about 0.1 L/s. Prior to sample collection the water column was exchanged 2.5 times. Pumped groundwater was collected in sterile 10 L Nalgene® containers in triplicates and transported back to the laboratory for immediate filtration. Sampling site 2 is a drinking water supply well in Entringen with a permanent pumping system, where a tap at the well head was used for sampling. Prior to sampling, the production well was operated for at least one hour to maintain steady state conditions. Samples were collected in triplicates in sterile 10 L Nalgene® containers, transported back to the laboratory and immediately filtered. In the laboratory, groundwaters were filtered sequentially through 8 µm (Millipore, TETP04700), 0.4 µm (Millipore, HTTP04700), and 0.2 µm (Millipore, GTTP04700) polycarbonate filters. The filters were frozen at −20 °C until further analysis. DNA was extracted from the 0.2 µm filters using the FastDNA spin kit (MP Biomedicals, Santa Ana, CA, USA) according to the manufacturer’s instructions.

#### Soil

Topsoil (0-10 cm depth) was collected using a sterile ethanol-washed spatula into sterile Corning™ Falcon 50mL Conical Tubes. Samples were transported at ambient temperature back to the laboratory (within ∼8 hours) and frozen at −80°C. DNA was extracted according to Lueders *et al*. (2004).

#### Sediment

Sediment was collected using a sterile ethanol-washed stainless steel corer with an inner diameter of 4 cm. Subsamples from 5 cm depth were removed carefully from the core using a sterile autoclaved stainless-steel spatula and placed into sterile Corning™ Falcon 50mL Conical Tubes before being frozen on dry ice immediately in the field. DNA was extracted from 0.5 g (wet weight) according to Lueders *et al*. (2004).

#### River water

River water samples were collected in triplicates into sterile 10 L Nalgene® canisters. Sample containers were held below the water surface (at ∼20 cm depth) and transported back to the laboratory for immediate filtering (i.e. within maximum 8 hours after retrieval) through a 0.2 µm polyethersulfone filter (Steritop; EMD Millipore). Filters were frozen at −20 °C until DNA extraction. DNA was extracted from the filters using the FastDNA spin kit (MP Biomedicals, Santa Ana, CA, USA) according to the manufacturer’s instructions.

Further details such as sample names, sampling coordinates, and sampling dates can be found in the supplementary information (Table S5).

### 16S rRNA gene amplicon sequencing

Bacterial and archaeal 16S rRNA genes were amplified using universal primers 515F: GTGYCAGCMGCCGCGGTAA (Parada, Needham et al. 2016) and 806R: GGACTACNVGGGTWTCTAAT (Apprill, McNally et al. 2015) fused to Illumina adapters. PCR mixtures for amplification contained (per 25 µl reaction): 0.5 µl of each primer (515F and 806R with Illumina tags; 10 µM stock concentration), 12.5 µl of 2x KAPA HiFi Hotstart Readymix (Kapa Biosystems, In., Wilmington, MA, USA), 0.5 µl BSA (10% stock solution), 10 µl of RNAse/DNAse-free water and 1 µl of template. The thermal profile used was: 3 min at 95 °C, 27 cycles of 95 °C 30 s, 55 °C 30 s, 72 °C 30 s and 5 min at 72 °C. Subsequent library preparation steps (Nextera, Illumina) and 250 bp paired-end sequencing with MiSeq (Illumina, San Diego, CA, USA) using v2 chemistry were performed by Microsynth AG (Balach, Switzerland) and between 40,000 to 132,000 read pairs per sample were obtained totaling to 2,368,742 read pairs with 1,166,187,315 nucleotides. Primers were removed from raw sequences and untrimmed sequences were discarded with Cutadapt v1.14 (Martin 2011) wrapped by Trim Galore! v0.4.5 (Felix Krueger, bioinformatics.babraham.ac.uk).

### 16S rRNA gene amplicon sequencing analysis software

The mock and environmental datasets were analyzed without (Extreme dataset) or with (all other datasets) primer trimming with Mothur, QIIME1, QIIME2, MEGAN as described below.

For Mothur analysis, Mothur v1.40.5 (Schloss, Westcott et al. 2009) was used with standard settings following the MiSeqSOP (Kozich, Westcott et al. 2013), except adjusting the cutoff of the reference alignment to the majority of aligned reads.

Briefly, paired-end sequences were merged and only those with maximum 8 homopolymers and maximum 275 bp were retained. SILVA v132 alignment was cut to the amplified region (position 11894-25319), and unique merged sequences were aligned to the cut SILVA alignment. The alignment region was refined (Balanced: position 1968-11546, all other: position 1968-11550) and only sequences aligned in that region were retained. Next, unique sequences were pre-clustered allowing for up to 2 nucleotide differences between sequences. Chimeras were removed by VSEARCH. Uncorrected pairwise distances were calculated and finally the sequences were clustered to OTUs at 0.03 (97% similarity) or 0.01 (99% similarity) cutoff and the consensus taxonomy for each OTU was retrieved.

For QIIME1 analysis, QIIME v1.9.1 was applied (Caporaso, Kuczynski et al. 2010) using fastq-join v1.3.1 (Aronesty 2013) for read merging, PyNAST v1.2.2 (Caporaso, Bittinger et al. 2010) for alignments, VSEARCH v2.3.4 (Rognes, Flouri et al. 2016) for OTU picking and chimera detection, uclust v1.2.22 (Edgar 2010) for taxonomy assignments with python v2.7.13 (van Rossum 1995) and matplotlib v1.4.3 (Hunter 2007).

For QIIME2 analysis, primer-free sequences were imported into QIIME2 q2cli v2018.06 (Bolyen, Rideout et al. 2019), visually inspected with demux (https://github.com/qiime2/q2-demux), and processed with DADA2 (Callahan, McMurdie et al. 2016) to remove PhiX contamination, trim reads, correct errors, merge read pairs and remove PCR chimeras, or merged with VSEARCH (Rognes, Flouri et al. 2016) followed by removal of PCR chimeras and Deblur (Amir, McDonald et al. 2017) to obtain representative ASV sequences. Representative sequences and their abundances were extracted by feature-table (McDonald, Clemente et al. 2012). A Naive Bayes classifier (Pedregosa, Varoquaux et al. 2011) was fitted with 16S rRNA gene sequences extracted from SILVA v132 (Quast, Pruesse et al. 2013) using the PCR primers of the investigated dataset. The representative sequences were classified by taxon using the fitted classifier (https://github.com/qiime2/q2-feature-classifier). QIIME2 plugins were executed with standard parameters, with DADA2 quality settings “--p-trunc-len-f” and “--p-trunc-len- r” for Extreme dataset 160 and 120 and for HMP and Balanced datasets with 200 and 120 or with Deblur parameter “--p-trim-length” 250 for Balanced dataset or 252 for Extreme and HMP datasets.

For MEGAN analysis, reads were merged using ClipAndMerge v1.7.4 (Peltzer, Jäger et al. 2016) and merged reads were aligned to SILVA using MALT v0.4.0 (Herbig, Maixner et al. 2016) with Parameters: --mode BlastN --alignmentType SemiGlobal -- sparseSAM. MEGAN v6.10.2 (Huson, Beier et al. 2016) assigned taxonomy (taxon path at genus level, all leaves) and abundances (counts) based on MALT alignments.

To facilitate reproducibility and to disseminate bioinformatics applications according to the FAIR principle (Wilkinson, Dumontier et al. 2016) all analysis software for the benchmarks was bundled in containers, using Singularity v2.4.1 (Kurtzer, Sochat et al. 2017) with Ubuntu 16.04.3 LTS and Conda/Bioconda 4.0.5 (ContinuumAnalytics, Inc., www.anaconda.com) and are publicly accessible. Information how to access these is available in the supplementary information (Table S6).

### Reference database

The SILVA v132 database (Quast, Pruesse et al. 2013) of 16S rRNA gene sequences, clustered at 99% similarity, was used as reference database. The used analysis software required specialized files that are indicated in the supplementary information (Table S7).

### Statistical analysis

The F-score was calculated as in (Kopylova, Navas-Molina et al. 2016):

Fscore = 2*precision*recall/(precision+recall)

where precision = (TP)/(TP+FP) and recall = (TP)/(TP+FN)

with TP = true positive, FP = false positive, FN = false negative

On the sequence level, only perfect matches and those with one mismatch to a reference sequence were counted as true positives. However, in the case where multiple ASVs/OTUs matched one reference sequence, only one was counted as expected and all others as unexpected (false positives).

### Jitter plot, Venn diagram, Upset plot, and Heatmap

All representative sequences were aligned to reference sequences with blastn v2.2.31+ and a jitter plot based on relative sequence abundances was produced by ggplot2 v2.2.1 (Wickham 2009) in R v3.4.4 (R Core Team 2018). The heatmap was done with pheatmap v1.0.8 (Kolde 2015), the upset plot with UpSetR v1.4.0 (Conway, Lex et al. 2017) and the venn diagram with gplots v3.0.1 (Warnes, Bolker et al. 2009) in R.

### Diversity indices and distances

Shannon index, Weighted UniFrac, Unweighted UniFrac and Bray-Curtis distance were calculated with the R-package phyloseq v1.22.3 (McMurdie and Holmes 2013) with the “estimate_richness” or “distance” function using ape v5.1 (Paradis, Claude et al. 2004). The Faith’s PD index was calculated with picante v1.7 (Kembel, Cowan et al. 2010) in R v3.4.4 (R Core Team 2018). For mock samples, expected alpha-diversity was calculated based on expected sequences and abundances. For environmental samples, Bray-Curtis distance or Unweighted UniFrac distances were subjected to NMDS (Non-metric Multidimensional Scaling) ordination and combined by Generalized Procrustes Analysis using plyr v1.8.4 (Wickham 2011) and FactoMineR v1.41 (Husson, Josse et al. 2008).

### Data availability

Raw sequencing data have been deposited at DDBJ/ENA/GenBank under BioProject accession number PRJNA563986 (https://www.ncbi.nlm.nih.gov/ bioproject/PRJNA563986).

## 5 Acknowledgements

We would like to thank Zhe Wang and Tillmann Lüders for the sediment sampling, Michael Lesch for soil sampling, and Franziska Schädler for DNA extraction. This work was funded by the Institutional Strategy of the University of Tübingen (German Research Foundation; DFG, ZUK 63) and by the Collaborative Research Center 1253 CAMPOS (Project 5: Fractured Aquifers), funded by the German Research Foundation (DFG, Grant Agreement SFB 1253/1 2017). S.K. is supported by an Emmy-Noether fellowship (DFG grant # 326028733) from the German Research Foundation. S.N. and A.P. acknowledge funding from the Deutsche

Forschungsgemeinschaft (core facilities initiative, KO-2313/6-1 and KO-2313-2, Institutional Strategy of the University of Tübingen, ZUK 63 as well as support from the project INF, SFB/TR 209 “Liver Cancer”). S.N. acknowledges funding from the Deutsche Forschungsgemeinschaft (DFG, German Research Foundation)- Project-ID 398967434 - TRR 261.

The authors acknowledge support by the state of Baden-Württemberg through bwHPC and the German Research Foundation (DFG) through grant no INST 37/935-1 FUGG (bwForCluster BinAC).

DS and SK designed the study in discussion with N.B., A.L.F. and S.N. D.S. analyzed and interpreted the data and D.S. (major part) and S.K. wrote the manuscript with the help of S.N. and N.B. Sampling and processing of environmental material was coordinated by N.B. and A.L.F. D.S. and A.P. wrote the analysis pipeline nf-core/ampliseq. All authors commented and approved the final manuscript. We declare that we have no conflicts of interest.

## References

Almeida, A., A. L. Mitchell, A. Tarkowska and R. D. Finn (2018). “Benchmarking taxonomic assignments based on 16S rRNA gene profiling of the microbiota from commonly sampled environments.” GigaScience 7(5): giy054.

Amir, A., D. McDonald, J. A. Navas-Molina, E. Kopylova, J. T. Morton, Z. Zech Xu, E. P. Kightley, L. R. Thompson, E. R. Hyde, A. Gonzalez and R. Knight (2017). “Deblur Rapidly Resolves Single-Nucleotide Community Sequence Patterns.” mSystems 2(2).

Antony-Babu, S., D. Stien, V. Eparvier, D. Parrot, S. Tomasi and M. T. Suzuki (2017). “Multiple Streptomyces species with distinct secondary metabolomes have identical 16S rRNA gene sequences.” Scientific Reports 7(1): 11089.

Apprill, A., S. McNally, R. Parsons and L. Weber (2015). “Minor revision to V4 region SSU rRNA 806R gene primer greatly increases detection of SAR11 bacterioplankton.” Aquatic Microbial Ecology 75.

Aronesty, E. (2013). ““Comparison of Sequencing Utility Programs”.” The Open Bioinformatics Journal 7: 1–8.

Bokulich, N. A., B. D. Kaehler, J. R. Rideout, M. Dillon, E. Bolyen, R. Knight, G. A. Huttley and J. G. Caporaso (2018). “Optimizing taxonomic classification of marker gene amplicon sequences.” PeerJ Preprints 6: e3208v3202.

Bokulich, N. A., S. Subramanian, J. J. Faith, D. Gevers, J. I. Gordon, R. Knight, D. A. Mills and J. G. Caporaso (2013). “Quality-filtering vastly improves diversity estimates from Illumina amplicon sequencing.” Nature methods 10(1): 57–59.

Bolyen, E., J. R. Rideout, M. R. Dillon, N. A. Bokulich, C. C. Abnet, G. A. Al-Ghalith, H. Alexander, E. J. Alm, M. Arumugam, F. Asnicar, Y. Bai, J. E. Bisanz, K. Bittinger, A. Brejnrod, C. J. Brislawn, C. T. Brown, B. J. Callahan, A. M. Caraballo-Rodríguez, J. Chase, E. K. Cope, R. Da Silva, C. Diener, P. C. Dorrestein, G. M. Douglas, D. M. Durall, C. Duvallet, C. F. Edwardson, M. Ernst, M. Estaki, J. Fouquier, J. M. Gauglitz, S. M. Gibbons, D. L. Gibson, A. Gonzalez, K. Gorlick, J. Guo, B. Hillmann, S. Holmes, H. Holste, C. Huttenhower, G. A. Huttley, S. Janssen, A. K. Jarmusch, L. Jiang, B. D. Kaehler, K. B. Kang, C. R. Keefe, P. Keim, S. T. Kelley, D. Knights, I. Koester, T. Kosciolek, J. Kreps, M. G. I. Langille, J. Lee, R. Ley, Y.-X. Liu, E. Loftfield, C. Lozupone, M. Maher, C. Marotz, B. D. Martin, D. McDonald, L. J. McIver, A. V. Melnik, J. L. Metcalf, S. C. Morgan, J. T. Morton, A. T. Naimey, J. A. Navas-Molina, L. F. Nothias, S. B. Orchanian, T. Pearson, S. L. Peoples, D. Petras, M. L. Preuss, E. Pruesse, L. B. Rasmussen, A. Rivers, M. S. Robeson, P. Rosenthal, N. Segata, M. Shaffer, A. Shiffer, R. Sinha, S. J. Song, J. R. Spear, A. D. Swafford, L. R. Thompson, P. J. Torres, P. Trinh, A. Tripathi, P. J. Turnbaugh, S. Ul-Hasan, J. J. J. van der Hooft, F. Vargas, Y. Vázquez-Baeza, E. Vogtmann, M. von Hippel, W. Walters, Y. Wan, M. Wang, J. Warren, K. C. Weber, C. H. D. Williamson, A. D. Willis, Z. Z. Xu, J. R. Zaneveld, Y. Zhang, Q. Zhu, R. Knight and J. G. Caporaso (2019). “Reproducible, interactive, scalable and extensible microbiome data science using QIIME 2.” Nature Biotechnology 37: 852–857.

Bray, J. R. and J. T. Curtis (1957). “An Ordination of the Upland Forest Communities of Southern Wisconsin.” Ecological Monographs 27(4): 325–349.

Callahan, B. J., P. J. McMurdie and S. P. Holmes (2017). “Exact sequence variants should replace operational taxonomic units in marker-gene data analysis.” The Isme Journal 11: 2639.

Callahan, B. J., P. J. McMurdie, M. J. Rosen, A. W. Han, A. J. A. Johnson and S. P. Holmes (2016). “DADA2: High resolution sample inference from Illumina amplicon data.” Nature methods 13(7): 581–583.

Callahan, B. J., J. Wong, C. Heiner, S. Oh, C. M. Theriot, A. S. Gulati, S. K. McGill and M. K. Dougherty (2019). “High-throughput amplicon sequencing of the full-length 16S rRNA gene with single-nucleotide resolution.” Nucleic Acids Research 47(18): e103.

Calus, S. T., U. Z. Ijaz and A. J. Pinto (2018). “NanoAmpli-Seq: a workflow for amplicon sequencing for mixed microbial communities on the nanopore sequencing platform.” GigaScience 7(12).

Caporaso, J. G., K. Bittinger, F. D. Bushman, T. Z. DeSantis, G. L. Andersen and R. Knight (2010). “PyNAST: a flexible tool for aligning sequences to a template alignment.” Bioinformatics 26(2): 266–267.

Caporaso, J. G., J. Kuczynski, J. Stombaugh, K. Bittinger, F. D. Bushman, E. K. Costello, N. Fierer, A. G. Peña, J. K. Goodrich, J. I. Gordon, G. A. Huttley, S. T. Kelley, D. Knights, J. E. Koenig, R. E. Ley, C. A. Lozupone, D. McDonald, B. D. Muegge, M. Pirrung, J. Reeder, J. R. Sevinsky, P. J. Turnbaugh, W. A. Walters, J. Widmann, T. Yatsunenko, J. Zaneveld and R. Knight (2010). “QIIME allows analysis of high-throughput community sequencing data.” Nature methods 7(5): 335–336.

Claesson, M. J., Q. Wang, O. O’Sullivan, R. Greene-Diniz, J. R. Cole, R. P. Ross and P. W. O’Toole (2010). “Comparison of two next-generation sequencing technologies for resolving highly complex microbiota composition using tandem variable 16S rRNA gene regions.” Nucleic Acids Research 38(22): e200–e200.

Conway, J. R., A. Lex and N. Gehlenborg (2017). “UpSetR: an R package for the visualization of intersecting sets and their properties.” Bioinformatics 33(18): 2938–2940.

Cuscó, A., C. Catozzi, J. Viñes, A. Sanchez and O. Francino (2018). “Microbiota profiling with long amplicons using Nanopore sequencing: full-length 16S rRNA gene and whole rrn operon.” F1000Research 7: 1755–1755.

D’Amore, R., U. Z. Ijaz, M. Schirmer, J. G. Kenny, R. Gregory, A. C. Darby, M. Shakya, M. Podar, C. Quince and N. Hall (2016). “A comprehensive benchmarking study of protocols and sequencing platforms for 16S rRNA community profiling.” BMC Genomics 17(1): 55.

de Muinck, E. J., P. Trosvik, G. D. Gilfillan, J. R. Hov and A. Y. M. Sundaram (2017). “A novel ultra high-throughput 16S rRNA gene amplicon sequencing library preparation method for the Illumina HiSeq platform.” Microbiome 5(1): 68.

de Voogd, N. J., D. F. R. Cleary, A. R. M. Polónia and N. C. M. Gomes (2015). “Bacterial community composition and predicted functional ecology of sponges, sediment and seawater from the thousand islands reef complex, West Java, Indonesia.” FEMS Microbiology Ecology 91(4).

DeSantis, T. Z., P. Hugenholtz, N. Larsen, M. Rojas, E. L. Brodie, K. Keller, T. Huber, D. Dalevi, P. Hu and G. L. Andersen (2006). “Greengenes, a Chimera-Checked 16S rRNA Gene Database and Workbench Compatible with ARB.” Applied and Environmental Microbiology 72(7): 5069.

Edgar, R. C. (2010). “Search and clustering orders of magnitude faster than BLAST.” Bioinformatics 26(19): 2460–2461.

Edgar, R. C. (2017). “Accuracy of microbial community diversity estimated by closed- and open-reference OTUs.” PeerJ 5: e3889.

Edgar, R. C. (2018). “Accuracy of taxonomy prediction for 16S rRNA and fungal ITS sequences.” PeerJ 6: e4652.

Ewels, P. A., A. Peltzer, S. Fillinger, J. Alneberg, H. Patel, A. Wilm, M. U. Garcia, P. D. Tommaso and S. Nahnsen (2019). “nf-core: Community curated bioinformatics pipelines.” bioRxiv: 610741.

Faith, D. (1992). “Conservation evaluation and phylogenetic diversity.” Biological Conservation 61(1): 1–10.

Hathaway, N., C. Parobek, J. Juliano and J. A Bailey (2017). “SeekDeep: Single-base resolution de novo clustering for amplicon deep sequencing.” Nucleic acids research 46(4): e21.

Head, I. M., J. R. Saunders and R. W. Pickup (1998). “Microbial Evolution, Diversity, and Ecology: A Decade of Ribosomal RNA Analysis of Uncultivated Microorganisms.” Microb Ecol 35(1): 1–21.

Herbig, A., F. Maixner, K. I. Bos, A. Zink, J. Krause and D. H. Huson (2016). “MALT: Fast alignment and analysis of metagenomic DNA sequence data applied to the Tyrolean Iceman.” bioRxiv.

Hugenholtz, P., B. M. Goebel and N. R. Pace (1998). “Impact of culture-independent studies on the emerging phylogenetic view of bacterial diversity.” Journal of bacteriology 180(18): 4765–4774.

Hunter, J. D. (2007). “Matplotlib: A 2D Graphics Environment.” Computing in Science & Engineering 9(3): 90–95.

Huson, D. H., A. F. Auch, J. Qi and S. C. Schuster (2007). “MEGAN analysis of metagenomic data.” Genome research 17(3): 377–386.

Huson, D. H., S. Beier, I. Flade, A. Górska, M. El-Hadidi, S. Mitra, H.-J. Ruscheweyh and R. Tappu (2016). “MEGAN Community Edition - Interactive Exploration and Analysis of Large-Scale Microbiome Sequencing Data.” PLOS Computational Biology 12(6): e1004957.

Huson, D. H., S. Mitra, H.-J. Ruscheweyh, N. Weber and S. C. Schuster (2011). “Integrative analysis of environmental sequences using MEGAN4.” Genome Research 21(9): 1552–1560.

Husson, F., J. Josse and S. Lê (2008). “FactoMineR: An R Package for Multivariate Analysis.” Journal of Statistical Software 25.

Jousset, A., C. Bienhold, A. Chatzinotas, L. Gallien, A. Gobet, V. Kurm, K. Küsel, M. C. Rillig, D. W. Rivett, J. F. Salles, M. G. A. van der Heijden, N. H. Youssef, X. Zhang, Z. Wei and W. H. G. Hol (2017). “Where less may be more: how the rare biosphere pulls ecosystems strings.” The ISME Journal 11(4): 853–862.

Jovel, J., J. Patterson, W. Wang, N. Hotte, S. O’Keefe, T. Mitchel, T. Perry, D. Kao, A. L. Mason, K. L. Madsen and G. K. S. Wong (2016). “Characterization of the Gut Microbiome Using 16S or Shotgun Metagenomics.” Frontiers in Microbiology 7: 459.

Kembel, S. W., P. D. Cowan, M. R. Helmus, W. K. Cornwell, H. Morlon, D. D. Ackerly, S. P. Blomberg and C. O. Webb (2010). “Picante: R tools for integrating phylogenies and ecology.” Bioinformatics 26(11): 1463–1464.

Kolde, R. (2015). pheatmap: Pretty Heatmaps. R package version 1.0.8.

Kopylova, E., J. A. Navas-Molina, C. Mercier, Z. Z. Xu, F. Mahé, Y. He, H.-W. Zhou, T. Rognes, J. G. Caporaso and R. Knight (2016). “Open-Source Sequence Clustering Methods Improve the State Of the Art.” mSystems 1(1): e00003–00015.

Kozich, J. J., S. L. Westcott, N. T. Baxter, S. K. Highlander and P. D. Schloss (2013). “Development of a Dual-Index Sequencing Strategy and Curation Pipeline for Analyzing Amplicon Sequence Data on the MiSeq Illumina Sequencing Platform.” Applied and Environmental Microbiology 79(17): 5112–5120.

Kurtzer, G. M., V. Sochat and M. W. Bauer (2017). “Singularity: Scientific containers for mobility of compute.” PLOS ONE 12(5): e0177459.

Laursen, M. F., M. D. Dalgaard and M. I. Bahl (2017). “Genomic GC-Content Affects the Accuracy of 16S rRNA Gene Sequencing Based Microbial Profiling due to PCR Bias.” Frontiers in microbiology 8: 1934–1934.

Lozupone, C. A., M. Hamady, S. T. Kelley and R. Knight (2007). “Quantitative and Qualitative β Diversity Measures Lead to Different Insights into Factors That Structure Microbial Communities.” Applied and Environmental Microbiology 73(5): 1576–1585.

Lueders, T., M. Manefield and M. W. Friedrich (2004). “Enhanced sensitivity of DNA- and rRNA-based stable isotope probing by fractionation and quantitative analysis of isopycnic centrifugation gradients.” Environmental Microbiology 6(1): 73–78.

Martin, M. (2011). “Cutadapt removes adapter sequences from high-throughput sequencing reads.” EMBnet.journal 17(1): 10–12.

McDonald, D., J. C. Clemente, J. Kuczynski, J. R. Rideout, J. Stombaugh, D. Wendel, A. Wilke, S. Huse, J. Hufnagle, F. Meyer, R. Knight and J. G. Caporaso (2012). “The Biological Observation Matrix (BIOM) format or: how I learned to stop worrying and love the ome-ome.” GigaScience 1(1): 1–6.

McMurdie, P. J. and S. Holmes (2013). “phyloseq: An R Package for Reproducible Interactive Analysis and Graphics of Microbiome Census Data.” PLOS ONE 8(4): e61217.

Mitra, S., M. Stärk and D. H. Huson (2011). “Analysis of 16S rRNA environmental sequences using MEGAN.” BMC Genomics 12(3): S17.

Muir, P., S. Li, S. Lou, D. Wang, D. J. Spakowicz, L. Salichos, J. Zhang, G. M. Weinstock, F. Isaacs, J. Rozowsky and M. Gerstein (2016). “The real cost of sequencing: scaling computation to keep pace with data generation.” Genome Biology 17: 53.

Musat, N., H. Halm, B. Winterholler, P. Hoppe, S. Peduzzi, F. Hillion, F. Horreard, R. Amann, B. B. Jørgensen and M. M. M. Kuypers (2008). “A single-cell view on the ecophysiology of anaerobic phototrophic bacteria.” Proceedings of the National Academy of Sciences of the United States of America 105(46): 17861–17866.

Nearing, J. T., G. M. Douglas, A. M. Comeau and M. G. I. Langille (2018). “Denoising the Denoisers: An independent evaluation of microbiome sequence error-correction methods.” PeerJ Preprints 6: e26566v26561.

Oliveira, C., L. Gunderman, C. A. Coles, J. Lochmann, M. Parks, E. Ballard, G. Glazko, Y. Rahmatallah, A. J. Tackett and D. J. Thomas (2017). “16S rRNA Gene-Based Metagenomic Analysis of Ozark Cave Bacteria.” Diversity 9(3): 31.

Parada, A. E., D. M. Needham and J. A. Fuhrman (2016). “Every base matters: assessing small subunit rRNA primers for marine microbiomes with mock communities, time series and global field samples.” Environmental Microbiology 18(5): 1403–1414.

Paradis, E., J. Claude and K. Strimmer (2004). “APE: Analyses of Phylogenetics and Evolution in R Language.” Bioinformatics (Oxford, England) 20: 289–290.

Pedregosa, F., G. Varoquaux, A. Gramfort, V. Michel, B. Thirion, O. Grisel, M. Blondel, P. Prettenhofer, R. Weiss, V. Dubourg, J. Vanderplas, A. Passos, D. Cournapeau, M. Brucher, M. Perrot and E. Duchesnay (2011). “Scikit-learn: Machine Learning in Python.” J. Mach. Learn. Res. 12: 2825–2830.

Peltzer, A., G. Jäger, A. Herbig, A. Seitz, C. Kniep, J. Krause and K. Nieselt (2016). “EAGER: efficient ancient genome reconstruction.” Genome Biology 17(1): 60.

Pester, M., N. Bittner, P. Deevong, M. Wagner and A. Loy (2010). “A ‘rare biosphere’ microorganism contributes to sulfate reduction in a peatland.” The ISME journal 4: 1591–1602.

Quast, C., E. Pruesse, P. Yilmaz, J. Gerken, T. Schweer, P. Yarza, J. Peplies and F. O. Glöckner (2013). “The SILVA ribosomal RNA gene database project: improved data processing and web-based tools.” Nucleic Acids Research 41(Database issue): D590–D596.

R Core Team (2018). “R: A language and environment for statistical computing. R Foundation for Statistical Computing, Vienna, Austria.”.

Rognes, T., T. Flouri, B. Nichols, C. Quince and F. Mahé (2016). “VSEARCH: a versatile open source tool for metagenomics.” PeerJ 4: e2584.

Schirmer, M., U. Z. Ijaz, R. D’Amore, N. Hall, W. T. Sloan and C. Quince (2015). “Insight into biases and sequencing errors for amplicon sequencing with the Illumina MiSeq platform.” Nucleic Acids Research 43(6): e37–e37.

Schloss, P. D., M. L. Jenior, C. C. Koumpouras, S. L. Westcott and S. K. Highlander (2016). “Sequencing 16S rRNA gene fragments using the PacBio SMRT DNA sequencing system.” PeerJ 4: e1869.

Schloss, P. D., S. L. Westcott, T. Ryabin, J. R. Hall, M. Hartmann, E. B. Hollister, R. A. Lesniewski, B. B. Oakley, D. H. Parks, C. J. Robinson, J. W. Sahl, B. Stres, G. G. Thallinger, D. J. Van Horn and C. F. Weber (2009). “Introducing mothur: Open-Source, Platform-Independent, Community-Supported Software for Describing and Comparing Microbial Communities.” Applied and Environmental Microbiology 75(23): 7537–7541.

Shannon, C. E. (1948). “A Mathematical Theory of Communication.” Bell System Technical Journal 27(3): 379–423.

Singer, E., B. Bushnell, D. Coleman-Derr, B. Bowman, R. M. Bowers, A. Levy, E. A. Gies, J.-F. Cheng, A. Copeland, H.-P. Klenk, S. J. Hallam, P. Hugenholtz, S. G. Tringe and T. Woyke (2016). “High-resolution phylogenetic microbial community profiling.” The Isme Journal 10: 2020.

Stoddard, S. F., B. J. Smith, R. Hein, B. R. K. Roller and T. M. Schmidt (2015). “rrnDB: improved tools for interpreting rRNA gene abundance in bacteria and archaea and a new foundation for future development.” Nucleic acids research 43(Database issue): D593–D598.

Straub, D. and A. Peltzer (2019). nf-core/ampliseq. Zenodo.

Taubert, M., C. Grob, A. Crombie, A. M. Howat, O. J. Burns, M. Weber, C. Lott, A.-K. Kaster, J. Vollmers, N. Jehmlich, M. von Bergen, Y. Chen and J. C. Murrell (2019). “Communal metabolism by Methylococcaceae and Methylophilaceae is driving rapid aerobic methane oxidation in sediments of a shallow seep near Elba, Italy.” Environmental Microbiology 0(0).

van Rossum, G. (1995). “Python Reference Manual.” CWI Report.

Větrovský, T. and P. Baldrian (2013). “The Variability of the 16S rRNA Gene in Bacterial Genomes and Its Consequences for Bacterial Community Analyses.” PLoS ONE 8(2): e57923.

Warnes, G. R., B. Bolker, L. Bonebakker, R. Gentleman, W. Huber, A. Liaw, T. Lumley, M. Maechler, A. Magnusson and S. Moeller (2009). “gplots: Various R programming tools for plotting data.” R package version 2(4): 1.

Wesolowska-Andersen, A., M. I. Bahl, V. Carvalho, K. Kristiansen, T. Sicheritz-Pontén, R. Gupta and T. R. Licht (2014). “Choice of bacterial DNA extraction method from fecal material influences community structure as evaluated by metagenomic analysis.” Microbiome 2(1): 19.

Wetterstrand, K. (2018). “DNA Sequencing Costs: Data from the NHGRI Genome Sequencing Program (GSP).” Retrieved 20-08-2018, 2018, from www.genome.gov/sequencingcostsdata.

Wickham, H. (2009). Ggplot2: Elegant Graphics for Data Analysis, Springer, New York, NY.

Wickham, H. (2011). “The Split-Apply-Combine Strategy for Data Analysis.” Journal of Statistical Software 40(1).

Wilkinson, M. D., M. Dumontier, I. J. Aalbersberg, G. Appleton, M. Axton, A. Baak, N. Blomberg, J.-W. Boiten, L. B. da Silva Santos, P. E. Bourne, J. Bouwman, A. J. Brookes, T. Clark, M. Crosas, I. Dillo, O. Dumon, S. Edmunds, C. T. Evelo, R. Finkers, A. Gonzalez-Beltran, A. J. G. Gray, P. Groth, C. Goble, J. S. Grethe, J. Heringa, P. A. C. ’t Hoen, R. Hooft, T. Kuhn, R. Kok, J. Kok, S. J. Lusher, M. E. Martone, A. Mons, A. L. Packer, B. Persson, P. Rocca-Serra, M. Roos, R. van Schaik, S.-A. Sansone, E. Schultes, T. Sengstag, T. Slater, G. Strawn, M. A. Swertz, M. Thompson, J. van der Lei, E. van Mulligen, J. Velterop, A. Waagmeester, P. Wittenburg, K. Wolstencroft, J. Zhao and B. Mons (2016). “The FAIR Guiding Principles for scientific data management and stewardship.” Scientific Data 3: 160018.

Yang, B., Y. Wang and P.-Y. Qian (2016). “Sensitivity and correlation of hypervariable regions in 16S rRNA genes in phylogenetic analysis.” BMC Bioinformatics 17(1): 135.

